# Phosphorylation-dependent conformational change in the Keratin 19 head domain: Implications for filament assembly and binding to GSK3β and 14-3-3

**DOI:** 10.64898/2026.11.06.764774

**Authors:** Youngheum Ji, Minchan Jeong, Jewoo Kim, Jiwoo Yeom, Min Ae Han, Chaerim Park, Sang-Hoon Lee, Eunmi Hong, Young Ho Ko, Pierre A. Coulombe, Jin Hae Kim, Chang-Hun Lee

## Abstract

Keratins are intermediate filament (IF) proteins expressed in various cell types, playing a crucial role in mechanical support and signaling pathways. Keratin 19 (K19) is predominantly found in gut epithelia and early embryonic ectoderm. K19 interacts with signaling proteins such as GSK3β and 14-3-3 to participate in cell signaling pathways. However, details on the binding sites of these proteins on K19, and specific molecular mechanisms governing these interactions, are lacking. Additionally, the N-terminal head domains of IFs including K19 are considered intrinsically disordered regions (IDRs), and their atomic-resolution structures are unavailable. Using NMR spectroscopy, we obtained the first solution structures of K19 head domain peptides. Structural models involving phosphomimetic peptides suggest a phosphorylation-dependent conformational change in the K19 head domain. The phosphomimetic S35E substitution alters the conformation of the head domain, affecting its association with keratin filaments and K19-binding partners. NMR spectral changes indicate that the S35E substitution affects the association of the K19 head domain to K8/K19 filaments and GSK3β, whereas no significant binding changes were observed with 14-3-3ζ and 14-3-3σ. Cell-based analyses further showed that the S35E mutation altered keratin filament organization and reduced the proximity between K19 and GSK3β in 293T cells, whereas no significant change was observed in the proximity between K19 and 14-3-3σ. Our findings provide insight into the structure-function relationship of the head domains in IF proteins with direct implications for the assembly of filaments and their role in signal modulation.

## Introduction

Intermediate filaments (IFs) are crucial components of the cytoskeleton in eukaryotic cells, providing mechanical support and structural stability, and supporting various cellular processes and pathways. Among the known 70 IF proteins, keratins play a particularly significant role in epithelial cells, where they form a dynamic network that enables cells to withstand and adapt to mechanical stress (*1*, *2*). A defining characteristic shared by all IF proteins, including keratins, is their tripartite domain structure comprising a central α-helical rod domain of fixed length flanked by non-helical head (N-terminal) and tail (C-terminal) domains. The rod domain, which is highly conserved across evolution, facilitates coiled-coil dimerization and IF assembly, whereas the head and tail domains, which are variable in length and substructure, regulate filament assembly and organization, mediate interactions with other cellular components (*1*, *2*), and are subject to post-translational modifications (*3*).

The head domains of keratin proteins are generally regarded as intrinsically disordered regions (IDRs), which lack a fixed three-dimensional conformation and remain highly flexible. These regions typically exhibit low complexity, consisting of repetitive sequences enriched in glycine and serine residues. A proposed explanation for the functional significance of keratin head domains is the glycine-loop hypothesis (*4*). This hypothesis suggests that glycine-rich loops within the head domains enhance filament flexibility and facilitate interactions with several signaling proteins, allowing cells to adapt to varying mechanical and signaling requirements. Phosphorylation of serine and threonine residues within the head domains of keratin proteins has also been implicated in regulating their dynamic behavior and interaction profiles (*5-8*). Supporting this idea, a study using ^13^C-labeled keratin head domains revealed an intense NMR peak, indicating dynamic motion upon dissociation from keratin filaments (*4*). Similarly, solid-state NMR studies and cryo-EM structural studies of other IF proteins, such as vimentin and desmin, demonstrated dynamic association and dissociation of their head domains (*9–12*). These findings suggest that IF head domains actively participate in intra- and intermolecular interactions. In essence, the head domain of IF proteins may function as key structural elements that regulate their mechanical properties and biological functions.

Given the dynamic and complex nature of keratin head domains, the divide-and-conquer approach that was originally devised to study IF rod domains (*10-16*) has also proven effective for characterizing smaller functional motifs within head domain regions. In this context, Eisenberg and colleagues used X-ray crystallography to identify a seven-amino-acid sequence (residues 58–64: SGMGSGG) in the K8 head domain that could adopt an amyloid-like conformation (*17*). This K8 sequence has been linked to Mallory-Denk body formation associated with chronic liver injury. Structural models incorporating alcohol molecules, along with analyses of the impact of three disease-associated variants (G191S, G225V, and G230C), provided atomic-resolution insight into the molecular mechanisms driving Mallory-Denk body formation. These findings underscore the intricate and dynamic nature of IF protein interactions and emphasize the importance of a detailed structural understanding for elucidating keratin-related pathological mechanisms (*17*).

Despite significant progress, the flexible and disordered nature of the head domains of several IF proteins poses a major challenge for structural characterization (*16*). Solution NMR spectroscopy has emerged as a powerful tool to study the structural and functional properties of disordered regions at atomic resolution (*18*, *19*). By preserving protein dynamics in aqueous samples during data acquisition, this method enables structural analysis under near-physiological conditions, making it particularly valuable for investigating intrinsically disordered proteins (IDPs) and IDRs (*20*). Among IF proteins, the head domain of keratin 19 (K19) is especially well-suited for solution NMR studies due to its simple repeat pattern and shorter length compared to other IF head domains.

K19, a type I intermediate filament protein, is distinct among keratins due to its small size. It is predominantly expressed in epithelial tissues, especially in simple epithelia and the embryonic ectoderm, and is frequently co-expressed with keratin 8 (K8), a major type II keratin found in the gut, liver and pancreas. K19 plays a vital role in preserving cellular architecture and resilience, notably in cells exposed to mechanical stress (*21*, *22*). In addition to its structural function, K19 contributes to cell differentiation, proliferation, and migration (*22*, *23*). Clinically, K19 expression serves as a biomarker for cancer detection (e.g., CYFRA 21-1 in non-small cell lung cancer), with elevated K19 levels frequently associated with epithelial-origin carcinomas (*24–26*). K19 interacts with a diverse array of cellular proteins, reinforcing its broad functional significance in cellular architecture, signaling, and cancer progression (*5*, *27*, *28*). As a type I IF protein, K19 forms heterodimers with type II keratins, particularly K8 but also K5, to establish a robust cytoskeletal framework. Beyond structural stabilization, K19 engages with proteins involved in cellular signaling and adhesion. For example, it associates with desmosome and hemidesmosome components, which are crucial for maintaining cell-cell and cell-matrix junctions (*29*, *30*). Additionally, K19 has been shown to bind 14-3-3, a family of adaptor proteins involved in a broad range of signal transduction pathways modulating cellular stress responses and cell death via apoptosis (6). In cancer cells, K19 also interacts with GSK3β, a key regulator of tumor cell proliferation, migration, and invasion, highlighting its significance in both normal epithelial homeostasis and oncogenesis (*5*). However, how phosphorylation-dependent changes in K19 affect filament structure and organization and its interactions with binding partners remains unclear.

In this study, we used NMR spectroscopy to determine the solution structure of overlapping peptides corresponding to specific segments of the K19 head domain. Structural modeling of a phosphomimetic variant of the head domain revealed a marked conformational shift compared to its wild-type (WT) counterpart. NMR titration experiments revealed interactions of the WT and phosphomimetic K19 head peptides with K8/K19 filaments, GSK3β, 14-3-3ζ, and 14-3-3σ. To further investigate the significance of site-specific phosphorylation events, we employed transmission electron microscopy (TEM) to examine the effects of the S35E substitution on filament assembly and structure. Additionally, co-sedimentation assays quantitatively confirmed that phosphorylation-mimicking variants, e.g., S35E, modulate the ability of K8/K19 filaments to bind key interacting partners. Cell-based analyses were performed to assess the effects of the S35E substitution on keratin filament organization and the physical proximity of K19 to GSK3β and 14-3-3σ. This study provides the first solution structure of the N-terminal segment (residues 6-38) of the head domain in an IF protein, K19, and elucidates the molecular mechanisms by which the K19 head domain interacts with keratin filaments and signaling proteins, GSK3β and 14-3-3.

## Results

### Solution NMR Structural Analysis of the Low-Complexity K19 Head Domain: A Divide-and-Conquer Approach

Low complexity is a defining and widespread feature of the amino acid sequence of the head and tail domains of intermediate filament proteins. This characteristic arises from repetitive sequences, notably glycine and serine repeats, which are conserved across various types of intermediate filament proteins, in particular, keratins. While such repetitive motifs enhance the flexibility and dynamic nature of the domains bearing them, they also introduce significant obstacles for structural investigations. Indeed, the inherent low complexity and dynamic properties of these regions complicate structural characterization using conventional techniques such as X-ray crystallography and NMR spectroscopy. In particular, NMR signal overlap, caused by recurring glycine and serine residues within a single polypeptide chain, makes it challenging to clearly distinguish individual signals, thereby reducing the precision and resolution of structure determinations. To overcome these limitations, we subdivided nearly the entire N-terminal half of the K19 head domain into distinct segments and synthesized short peptides corresponding to these regions (Fig. 1A). Conversely, the C-terminal half of the K19 head domain (G39–N79) consists almost entirely of repetitive serine and glycine residues, making it unsuitable for further analysis due to major signal overlap in NMR spectra. Consequently, we excluded this region from further study.

**Fig. 1.**
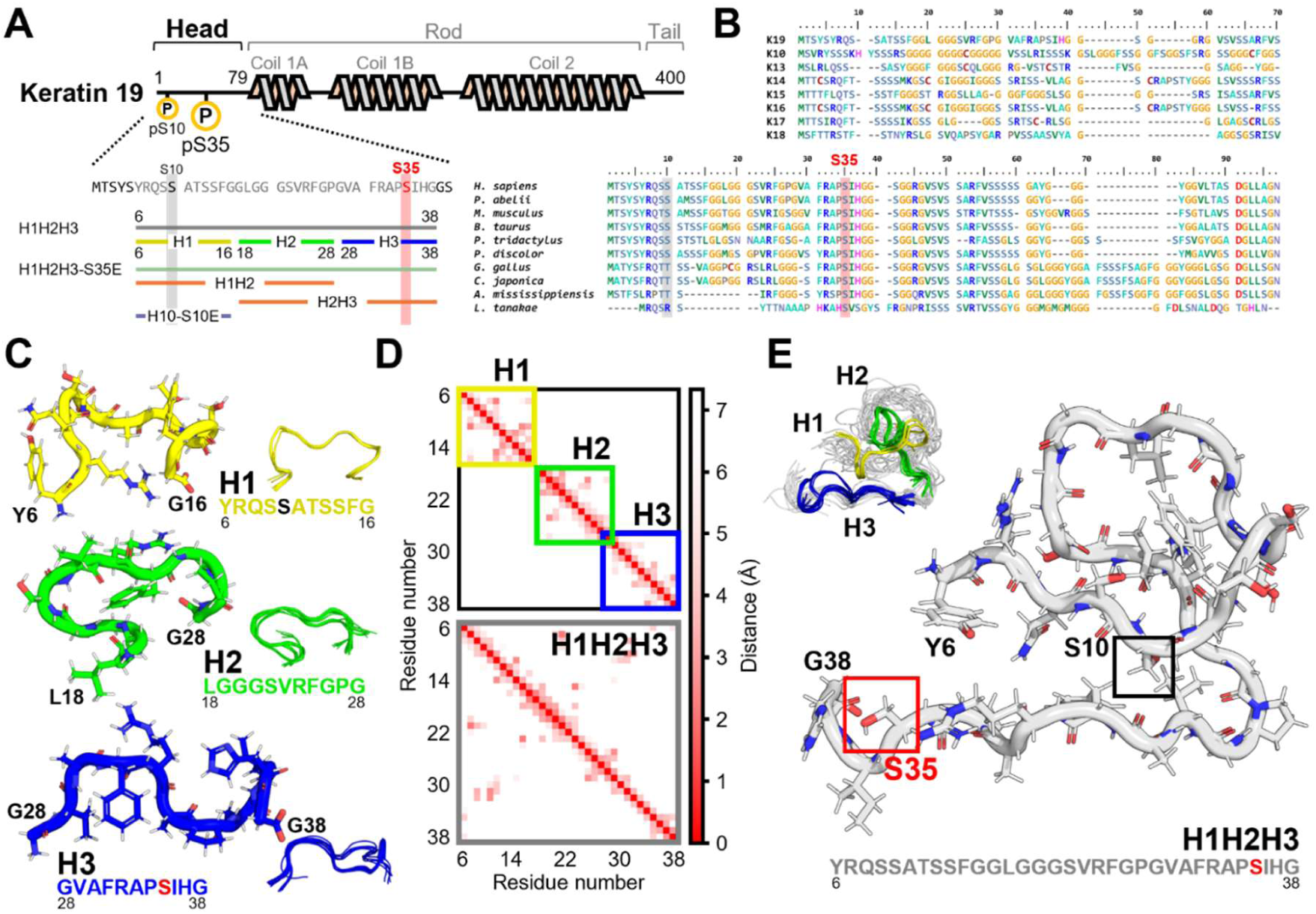
Amino acid composition and structural distinctions in the K19 head domain. **(A)** The sequence of the K19 head domain is displayed, with key phosphorylation sites S10 and S35 marked in red. The color-coded sequence strips align with the structural elements illustrated below. **(B)** Multiple sequence alignments of Type I keratin head domains, including interspecies comparisons, are provided. The alignment focuses on the anterior half of the K19 head domain, covering phosphorylation sites S10 and S35, which are highly conserved across species, emphasizing their potential functional importance. **(C)** The NMR-derived structural model depicts three short regions (H1: yellow, H2: green, H3: blue) with their superimposed structures, illustrating their relative spatial arrangement and conformational interplay. Corresponding PDB IDs are H1 (8ZG2), H2 (8ZG3), and H3 (8Y3S). **(D)** A comparative analysis of the merged contact map between H1, H2, and H3 versus the contact map of the full H1H2H3 region is shown. This analysis spans residues Y6 to G38, with color intensity representing interaction strength between residues. Notably, the full H1H2H3 contact map reveals long-range contacts absent in the merged individual maps, suggesting a cooperative folding pattern unique to the intact structure. **(E)** The computed H1H2H3 structural model displays a distinct three-dimensional conformation, which differs significantly from the isolated short peptide structures. This underscores the necessity of analyzing the complete H1H2H3 assembly to identify folding interactions that are not evident in individual segment models.

The three peptides, H1 (Y6-G16, PDB ID 8ZG2), H2 (L18-G28, PDB ID 8ZG3), and H3 (G28-G38, PDB ID 8Y3S) (Fig. 1C), were designed to serve as localized structural representations of the K19 head domain. While each peptide yielded a locally defined NMR-derived conformation individually, it remains unclear whether they function as discrete structural units within the full-length domain. To investigate this, we expanded our analysis to the longer segment, represented by H1H2H3, to assess its three-dimensional folding behavior. This experimental approach aligns with the “divide-and-conquer strategy” frequently applied in solution NMR to mitigate signal overlap and facilitate the characterization of structurally dynamic regions (*31*). To bridge intermediate regions, longer peptides H1H2 (Y6-G28, PDB ID 8ZUG) and H2H3 (L18-G38, PDB ID 8XTO) were synthesized and analyzed as well (Fig. S1). A structural model of the full-length construct, H1H2H3 (residues Y6-G38, PDB ID 8XTN), was determined with a backbone root-mean-square deviation (RMSD) of 2.5 Å across the top 20 energy-stable models. Notably, H1H2H3 adopted a distinct three-dimensional conformation that differed markedly from the linear or independently folded arrangements observed in the shorter peptides (Fig. 1E). This finding underscores that low-complexity domains like the K19 head domain rely on intricate folding interactions that cannot be inferred from isolated segments alone.

Moreover, the short keratin head domain segments may not independently adopt the local structures predicted by the ‘glycine-loop’ hypothesis (*4*). However, the short peptides in this study were instrumental in aiding the structural analysis of H1H2H3 by providing critical reference data to resolve signal overlap, particularly in samples lacking isotope enrichment. The divide-and-conquer methodology using solution NMR thus underscores the benefit of analyzing short peptides and full-length constructs in parallel when determining the structural organization of low-complexity domains such as the K19 head domain.

### Conformational Impact of S35 Phosphorylation in the K19 Head Domain

The potential significance of the two serine phosphorylation sites, S10 and S35, was initially assessed through multiple sequence alignment across species (Fig. 1B). Utilizing Clustal Omega, we examined sequences from ten species, including humans, reptiles, and fish. The alignment was conducted using default parameters (gap opening penalty of 10, gap extension penalty of 0.1, and BLOSUM62 substitution matrix). The analysis revealed that both S10 and S35 exhibit strong conservation across all species assessed, emphasizing their potential functional relevance (*32*). Expanding on these findings, a sequence-based phosphopeptide binding prediction was performed for the serine sites within the K19 head domain, a possible interaction site for 14-3-3 proteins (Table S3) (*33*). Among the serine residues analyzed in H1H2H3, S35 was the only site to yield scores surpassing the predefined cut-off threshold, reinforcing its likely functional importance. In contrast, structural comparison of the phosphomimetic H1-S10E peptide with H1 revealed only modest conformational changes (Fig. S2). Also, cell-based experiments and peptide mapping identified S35 as a major phosphorylation site (*7*). Based on these findings, S35 was prioritized for follow-up studies of phosphomimetic mutants.

To assess the structural consequences of serine phosphorylation within the K19 head domain, we engineered a phosphomimetic mutant peptide, H1H2H3-S35E, by replacing serine at position 35 with glutamic acid. This substitution emulates the negative charge introduced by phosphorylation, allowing us to investigate its impact on peptide folding and residue interactions. NMR-based structural analyses revealed conformational differences between the WT H1H2H3 peptide and its S35E mutant (Fig. 2A). Unlike WT H1H2H3, which exhibits contact between the N-terminal region and the C-terminal region, the S35E mutant shows a significant conformational shift and loses these interactions. This shift is accompanied by a rearrangement of the aromatic ring of F31, which likely contributes to the observed structural reorganization. The contact map further conveys that the S35E mutation disrupts key long-range interactions present in WT H1H2H3, particularly those involving Q8, S10, F31, and R32 (Fig. 2D). Loss of these interactions may contribute to the altered structural flexibility of the mutant peptide. Chemical shift perturbation (CSP) analysis (Fig. 2C) revealed that, beyond the expected perturbations around S35 due to its replacement by E35, signal disappearance occurs around S10 and S13, alongside substantial signal shifts at P34-G38, indicating localized structural rearrangements. These findings align with the observed disruption of long-range residue-residue interactions caused by the S35E substitution. Collectively, these results suggest that the phosphomimetic S35E substitution promotes structural rearrangement within the K19 head domain, potentially playing a crucial role in modulating both its conformational stability and functional dynamics.

**Fig. 2.**
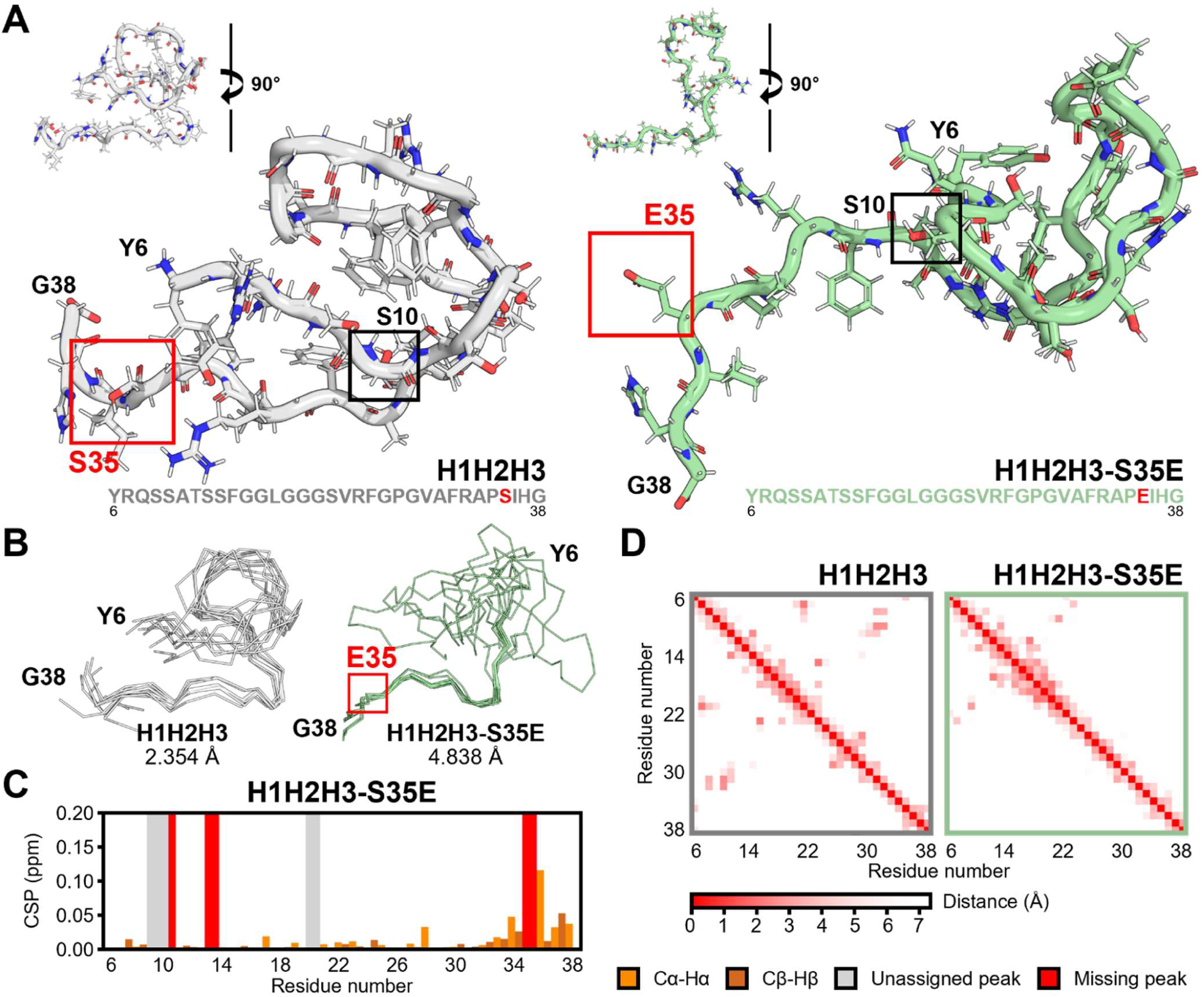
Structural variations between the WT H1H2H3 model and the phosphomimetic mutant H1H2H3-S35E. **(A)** NMR-derived structural models of H1H2H3 and H1H2H3-S35E, presented with orthogonal views rotated by 90 degrees for direct comparison. The S35E substitution disrupts the interaction between the N-terminal and the C-terminal region and exposes the C-terminal region to solvent, leading to significant alterations in the overall three-dimensional conformation. **(B)** Superimposition of the 10 lowest-energy NMR-derived structural models of H1H2H3 and H1H2H3-S35E. The RMSD values were calculated over all residues. **(C)** Chemical shift perturbation (CSP) plot for H1H2H3 and H1H2H3-S35E reveals substantial differences in chemical shifts following the S35E substitution. The CSP plot quantifies perturbations in the backbone chemical shifts (Cα-Hα, orange; Cβ-Hβ, brown), highlighting signal changes between H1H2H3 and H1H2H3-S35E. In this plot, gray and red bars correspond to unassigned (not observed in either) and missing (observed in H1H2H3, but absent in H1H2H3-S35E) peaks, respectively. **(D)** The contact maps illustrate patterns of residue interactions, where dot intensity correlates with the proximity of Cβ atoms, with closer distances denoted by darker red hues.

### Exploring a Role for K19 S35 Phosphorylation in the Assembly and Structure of K8-K19 Filaments

Building upon the impact of phosphorylation at S35, we subsequently investigated how the S35 site in the K19 head domain contributes to keratin filament assembly (Fig. 3). Previous studies showed that cells transfected with the K19-S35A mutant display collapsed or shorter keratin networks compared with the cells expressing wild-type K19 or an S10A variant (*7*). We hypothesized that phosphorylation at S35 might modulate the processes of filament assembly in keratin filaments. To test this hypothesis, we employed 2D ^1^H-^13^C heteronuclear single quantum coherence (HSQC) NMR spectroscopy to monitor interactions involving S35, TEM imaging to visualize filament morphology, sedimentation assays to assess filament assembly quantitatively, and cell-based transfection assays to examine keratin filament organization in 293T cells.

**Fig. 3.**
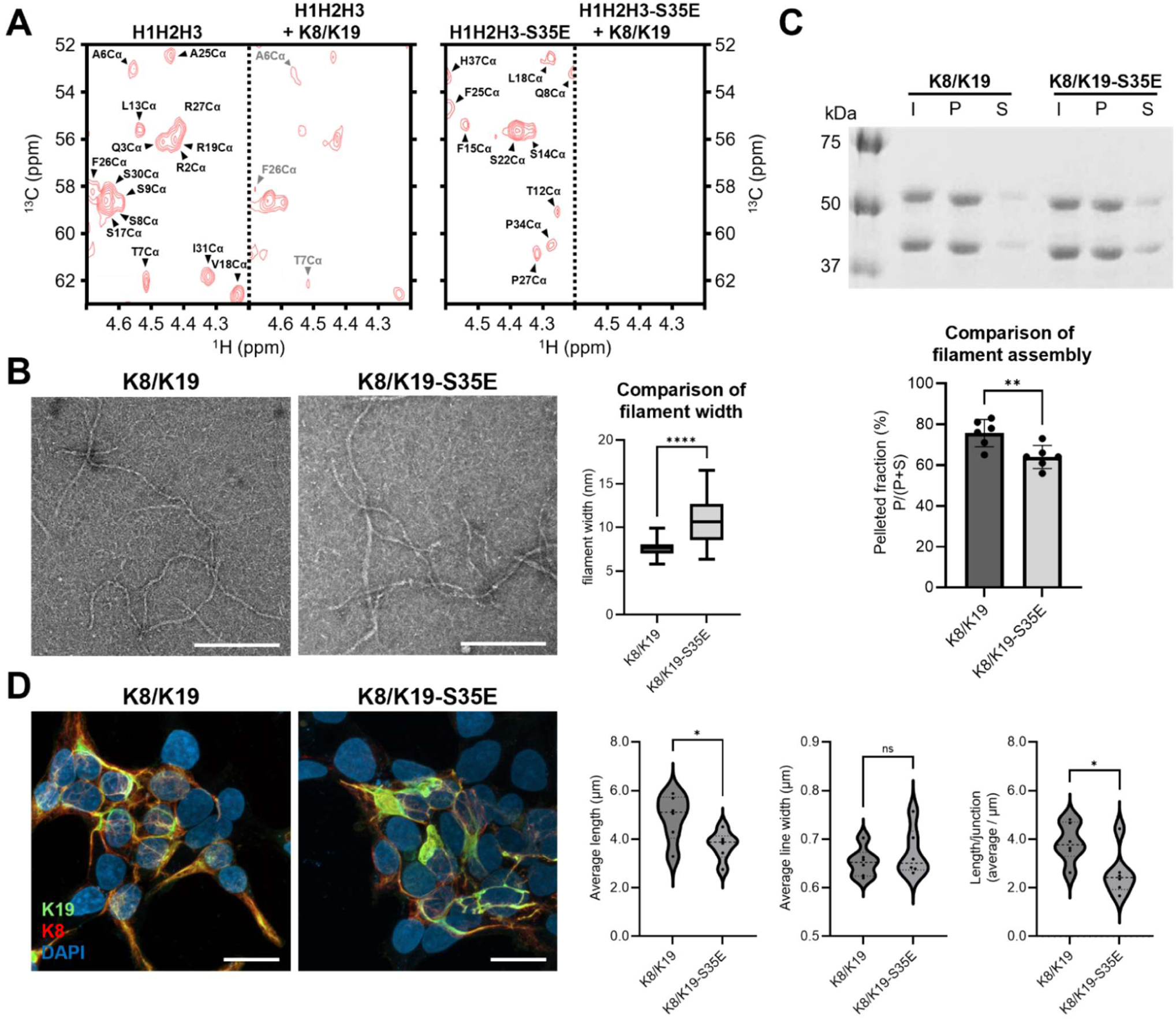
Impact of the S35 residue in the K19 head domain on K8/K19 filament structure and filament assembly. **(A)** 2D ^1^H-^13^C HSQC NMR strips comparing H1H2H3 and H1H2H3-S35E before and after the addition of K8/K19 filaments showed complete signal broadening of H1H2H3-S35E upon addition of K8/K19 filaments, indicating enhanced association of H1H2H3-S35E with K8/K19 filaments. **(B)** TEM imaging of K8/K19 filaments and K8/K19-S35E mutant filaments revealed a 42.1% increase in average filament width, from 7.55 nm (K8/K19) to 10.73 nm (K8/K19-S35E), based on measurements of *n* = 100 filaments. Scale bars, 200 nm. **(C)** High-speed sedimentation assays showed a significantly reduced pelleted fraction in K8/K19-S35E compared with K8/K19, indicating altered filament assembly associated with the S35E substitution. I, input; P, pellet; S, supernatant. The number of independent experiments was as follows: K8/K19 (*n* = 6), K8/K19-S35E (*n* = 6). **(D)** Representative confocal immunofluorescence images of 293T cells co-transfected with K8 and either wild-type K19 or K19-S35E. K19, green; K8, red; DAPI, blue. Quantification of keratin filament morphology by ridge detection showed significant reductions in average filament length and length per junction in K8/K19-S35E compared with K8/K19, whereas no significant difference was observed in average line width. Scale bars, 20 μm. Statistical significance was determined using Welch’s t-test for (B) and (C) and an unpaired Student’s t-test for (D). \**P* < 0.05, \*\**P* < 0.01, \*\*\*\**P* < 0.0001; ns, not significant.

The 2D NMR spectra of H1H2H3 and H1H2H3-S35E exhibit marked signal broadening upon addition of K8/K19 filaments, which reflects formation of high-molecular-weight peptide-filament complexes (Fig. 3A). The S35E peptide exhibited more pronounced signal broadening and missing peaks than that of H1H2H3, as evidenced by the complete disappearance of signals upon addition of K8/K19 filaments. These observations suggest that phosphorylation at S35 enhances the association of the K19 head domain into the K8/K19 filament. TEM imaging (Fig. 3B) reveals notable morphological differences between K8/K19 and K8/K19-S35E mutant filaments. Quantitative analysis showed that the S35E mutant filaments have a significantly increased width (10.73 nm ± 2.534 nm) compared to WT filaments (7.55 nm ± 0.813 nm; *P* < 0.0001), indicating substantial structural alterations driven by phosphorylation at S35. Notably, TEM images reveal regions of partial unraveling and irregular thickening along the mutant filaments, suggesting a distinct filament architecture. These observations are further corroborated by sedimentation assays (Fig. 3C), which demonstrate a significant reduction in the assembly efficiency of mutant filaments (64.00%) relative to WT filaments (75.67%; *P* = 0.0084). To assess whether these structural alterations were also observed in cells, we examined keratin filament organization in 293T cells co-transfected with K8 and either WT K19 or K19-S35E (Fig. 3D). Quantitative ridge-detection analysis showed significant reductions in average filament length and length per junction (the average filament segment length between junctions) in K8/K19-S35E cells compared with K8/K19 cells, whereas no significant difference was observed in average line width. Collectively, these findings suggest that the phosphomimetic S35E substitution affects filament assembly, stability and width of K8-K19 filaments, underscoring its significant contribution in maintaining the structural and functional integrity of K8/K19 filaments.

### Exploring a Role for K19 S35 Phosphorylation in Regulating Binding to GSK3β

We next explored how the structural changes induced by S35 phosphorylation on K19 influence its binding interactions with GSK3β, a key signaling molecule whose binding is reportedly markedly affected by this modification (*5*). GSK3β has been implicated in cancer and serves as a pivotal regulator of tumor progression by modulating processes such as cell proliferation, apoptosis, and metastasis. Previous studies have suggested that phosphorylation at S35 in K19 may play a crucial role in regulating GSK3β, offering a potential mechanism through which phosphorylation could impact signaling pathways linked to cancer progression (*5*). Gaining a deeper understanding of these interactions is essential for uncovering the functional role of phosphorylation at S35 in regulating binding partners, which could ultimately affect downstream signaling networks.

NMR binding assays revealed notable differences in the binding profiles of GSK3β to the WT H1H2H3 and the phosphomimetic mutant H1H2H3-S35E. Clear differences occur in the 2D NMR spectra, as manifested via peak shifts, peak disappearance, and changes in intensity (Fig. 4A). Noticeable peak shifts and signal broadening occur when mixing WT H1H2H3 and GSK3β. By contrast, the less pronounced spectral perturbations were observed with H1H2H3-S35E upon addition of GSK3β. This suggests that phosphorylation at S35 reduces the binding affinity between H1H2H3 and GSK3β by inducing conformational changes. To ensure that the GST tag did not affect the binding assays, control experiments were performed using H1H2H3 + GST and H1H2H3-S35E + GST. These controls confirmed that the binding interactions observed in the presence of GSK3β are independent of the GST tag. CSP analysis (Fig. 4B) further clarified the structural basis of these functional alterations. Addition of GSK3β to H1H2H3 results in marked chemical shift perturbations, particularly around F31-H37, indicating strong interactions. When using H1H2H3-S35E, these perturbations are attenuated, suggesting a decreased affinity to GSK3β.

**Fig. 4.**
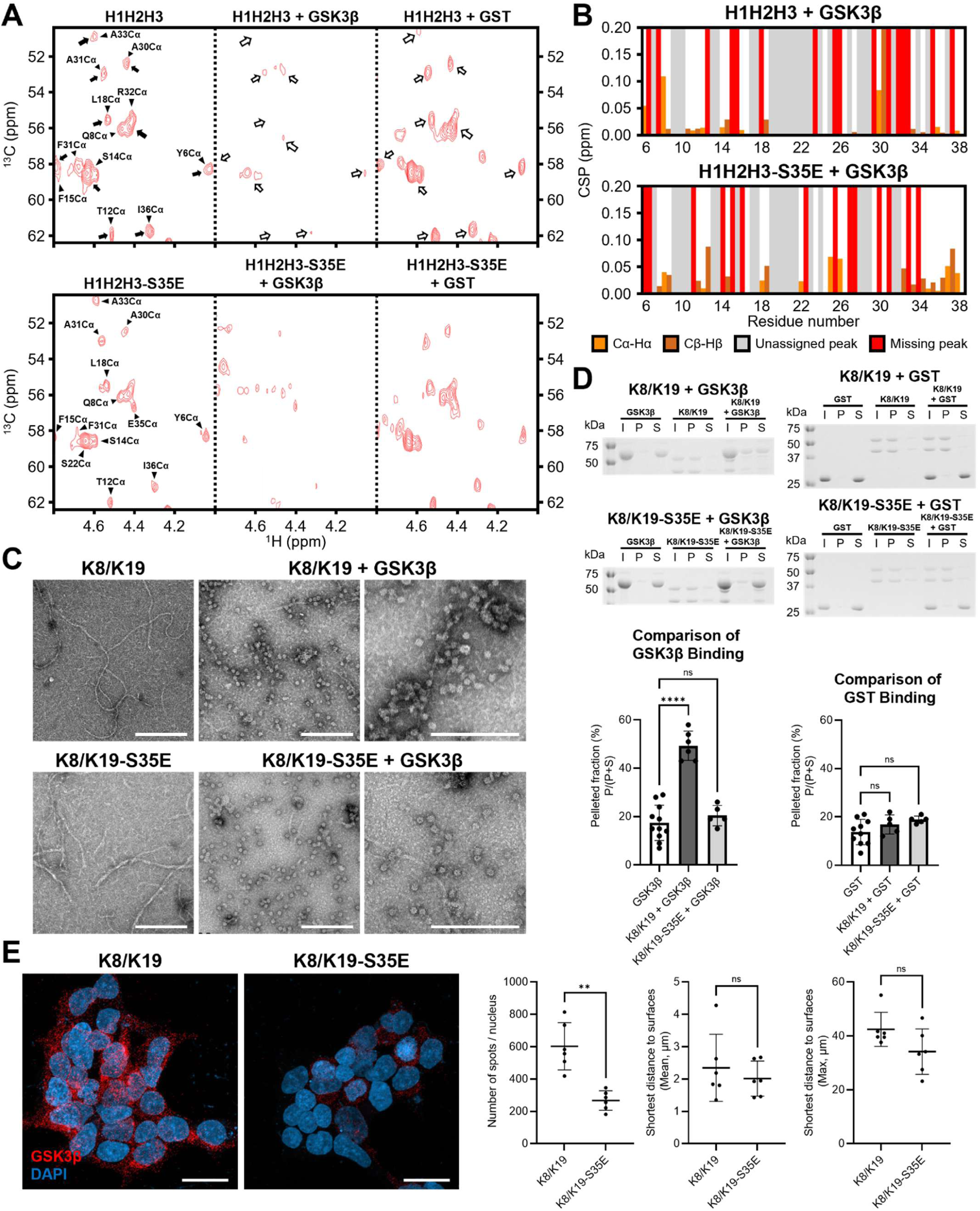
Comparison of GSK3β association with WT and S35E. **(A)** 2D ^1^H-^13^C HSQC NMR spectra demonstrating the impact of GSK3β binding on H1H2H3 and H1H2H3-S35E. The Cα region is shown, with GST included as a control for potential effects of the GST tag. Regions affected by GSK3β addition are indicated by arrows. **(B)** CSP plots comparing H1H2H3 + GSK3β and H1H2H3-S35E + GSK3β. CSPs for Cα–Hα and Cβ–Hβ correlations are shown in orange and brown, respectively. Gray and red bars indicate unassigned and missing peaks, respectively. **(C)** TEM images of K8/K19 filaments and K8/K19-S35E mutant filaments in the presence of GSK3β. Scale bars, 200 nm. **(D)** Co-sedimentation assays of GSK3β with K8/K19 and K8/K19-S35E filaments. Band intensities were quantified using ImageJ, and relative binding was expressed as the pelleted fraction, P/(P + S). The number of independent experiments was as follows: GSK3β (*n* = 11), K8/K19 + GSK3β (*n* = 6), K8/K19-S35E + GSK3β (*n* = 5), GST (*n* = 10), K8/K19 + GST (*n* = 5), and K8/K19-S35E + GST (*n* = 5). **(E)** Representative confocal images of a proximity ligation assay (PLA) detecting the in situ interaction between GSK3β and K19 in GSK3β-overexpressing 293T cells co-transfected with K8 and either wild-type K19 or K19-S35E. Red puncta represent positive PLA signals, and nuclei were counterstained with DAPI (blue). Quantification of PLA signals showed a significant reduction in the number of spots per nucleus in K8/K19-S35E compared with K8/K19, whereas no significant differences were observed in the mean or maximum distances of PLA spots from the nuclear surfaces. Scale bars, 20 μm. Statistical significance was determined using one-way ANOVA followed by Dunnett’s post hoc test for (D) and an unpaired Student’s t-test for (E). \*\**P* < 0.01, \*\*\*\**P* < 0.0001; ns, not significant.

TEM imaging (Fig. 4C) provided further support for the effects of S35E substitution. In the K8/K19 filament + GSK3β condition, evidence of binding of GSK3β can be seen at regular intervals along K8/K19 filaments, indicating stable, specific interactions. These “beads-on-a-string” structures, with beads ∼16 nm in diameter, are consistent with the expected size of GST-fused GSK3β dimer or oligomer, as reported in a previous X-ray crystallographic study (PDB ID 1I09) (*34*). This observation suggests that the bead-like structures observed by TEM correspond to GST-fused GSK3β associated with K8/K19 filaments. In contrast, evidence of binding interactions is markedly reduced in the K8/K19-S35E mutant filaments + GSK3β condition, suggesting that S35 phosphorylation weakens the interaction between the K19 head domain and GSK3β. Control experiments using GST alone confirmed that GST alone did not account for the observed association, as no specific binding occurs in the absence of GSK3β. This observation aligns with the reduced binding affinity observed in the NMR assays, further demonstrating that S35 phosphorylation alters the GSK3β-K19 head domain interaction.

Co-sedimentation assays were next performed to quantitatively assess the effect of phosphorylation on GSK3β binding. Pre-assembled K8/K19 filaments (WT or S35E) were incubated with GSK3β and subjected to ultracentrifugation to separate the pellet and supernatant fractions. The proportion of GSK3β bound to the filaments was quantified via densitometric analysis of Coomassie-stained SDS-PAGE gels using the ImageJ software. GSK3β binds significantly more to WT filaments (49.33%) compared to controls without filaments (17.45%; *P* < 0.0001), a 2.8-fold increase. By contrast, GSK3β binding to S35E mutant filaments (20.40%) does not differ significantly from controls (*P* = 0.6300). These results indicate that phosphorylation at K19 S35 greatly reduces GSK3β binding to K8/K19 filaments. During purification, GSK3β appeared as two distinct bands, identified as intact GSK3β by LC-MS and western blot analysis (Fig. S4) and consistent with established approaches for analysis of this kinase (*35*). To verify that these results were specific to GSK3β and not due to GST tag artifacts, parallel assays were conducted using GST alone. No significant differences were observed among GST alone (13.70%), GST with WT filaments (16.80%; *P* = 0.3556), or GST with S35E filaments (18.80%; *P* = 0.0855), ruling out GST-related artifacts.

To determine whether the S35E-dependent difference in the K19–GSK3β interaction observed in vitro was also evident in cells, we performed a proximity ligation assay (PLA) in GSK3β-overexpressing 293T cells (Fig. 4E). The number of PLA spots per nucleus was significantly reduced in cells expressing K19-S35E compared with WT K19, whereas no significant differences were observed in the mean or maximum distances of PLA spots from the nuclear surfaces. These results indicate reduced spatial proximity between K19-S35E and GSK3β compared with WT K19 in transfected 293T cells.

### Exploring a Role for K19 S35 Phosphorylation in Regulating Binding to 14-3-3ζ and 14-3-3σ

We next evaluated the binding interaction between the head domain of K19 and 14-3-3 proteins. Comparisons of the ^1^H-^13^C HSQC spectra of H1H2H3 and H1H2H3-S35E, both before and after the addition of 14-3-3ζ and 14-3-3σ, reveal binding of both peptides to the 14-3-3 proteins. The data suggest that 14-3-3 binding to K8-K19 is considerably weaker than that of GSK3β since modest (yet reproducible) differences in the perturbation pattern occur between the peptides and these two 14-3-3 proteins (Fig. 5A). Quantitative comparison of K19 head peptide binding to 14-3-3 proteins was precluded by NMR signal broadening and overlap with 14-3-3 resonances (Fig. 5A).

**Fig. 5.**
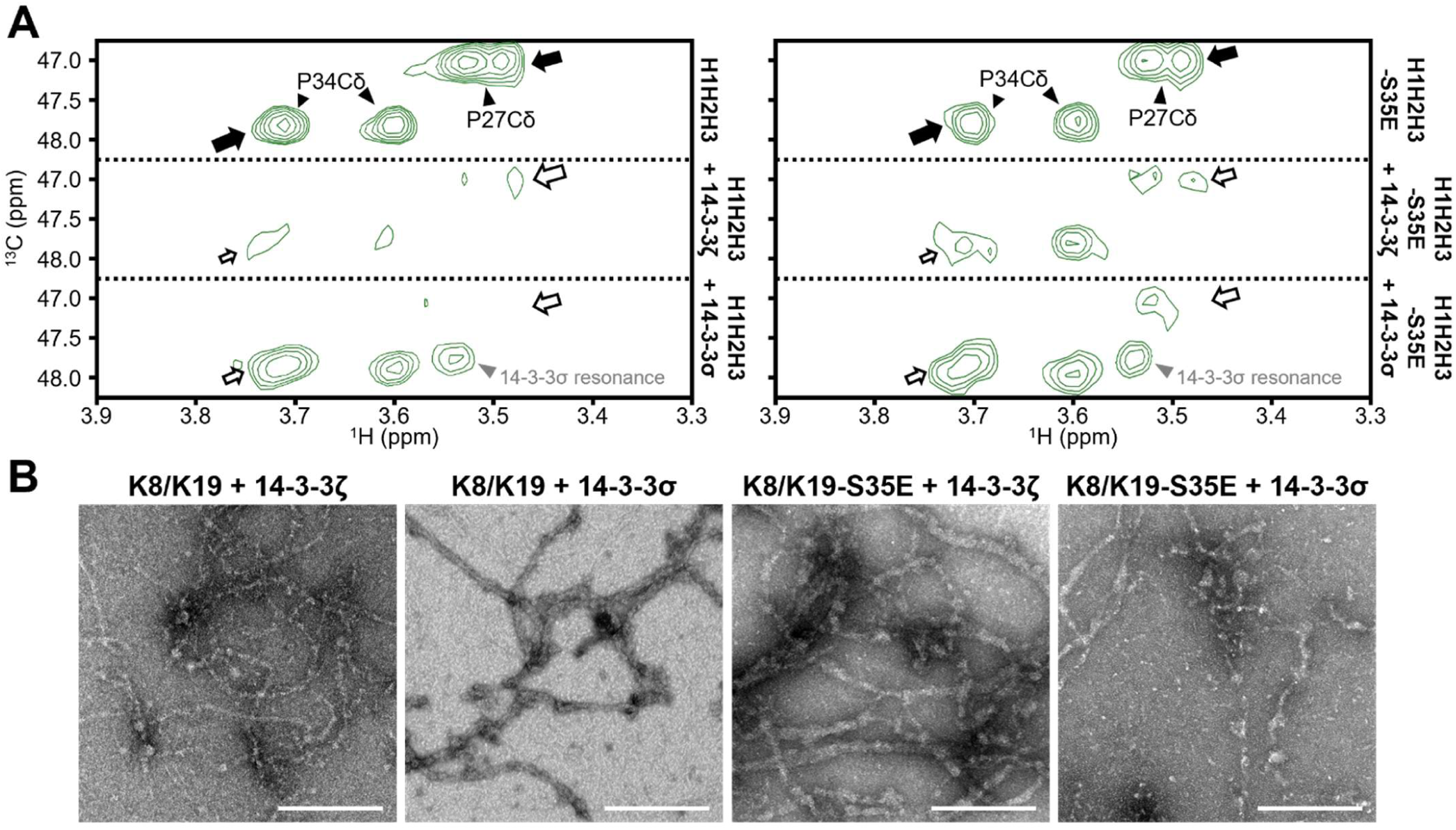
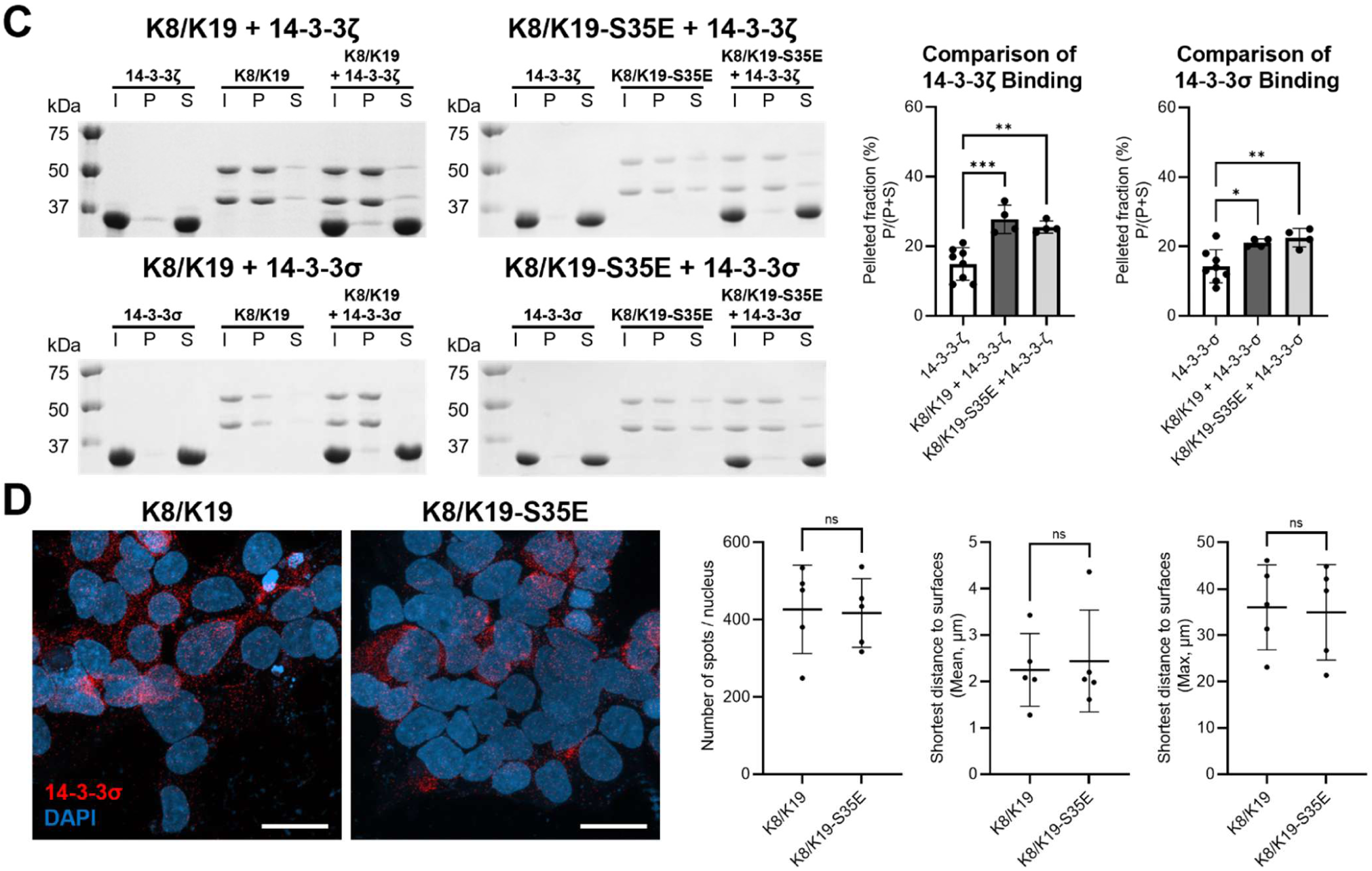
Comparison of the binding of 14-3-3ζ and 14-3-3σ to K19 as a function of S35 phosphorylation. **(A)** 2D ^1^H-^13^C HSQC NMR spectra illustrating the effects of 14-3-3ζ and 14-3-3σ interactions with H1H2H3 and H1H2H3-S35E. The focus is on the proline Cδ region. Signals affected by the addition of 14-3-3 are indicated by arrows. **(B)** TEM images depicting K8/K19 filaments and K8/K19-S35E mutant filaments in the presence of 14-3-3ζ and 14-3-3σ. 14-3-3ζ and 14-3-3σ were purified without GST tags. Scale bars, 200 nm. **(C)** Co-sedimentation assays comparing the binding of 14-3-3ζ and 14-3-3σ to K8/K19 and K8/K19-S35E filaments. The number of independent experiments was as follows: 14-3-3ζ (*n* = 8), K8/K19 + 14-3-3ζ (*n* = 4), K8/K19-S35E + 14-3-3ζ (*n* = 4), 14-3-3σ (*n* = 8), K8/K19 + 14-3-3σ (*n* = 4), and K8/K19-S35E + 14-3-3σ (*n* = 4). **(D)** Representative confocal images of a PLA detecting the in situ interaction between 14-3-3σ and K19 in 14-3-3σ-overexpressing 293T cells co-transfected with K8 and either wild-type K19 or K19-S35E. Red puncta represent positive PLA signals, and nuclei were counterstained with DAPI (blue). Quantification of PLA signals showed no significant differences in the number of spots per nucleus or in the mean and maximum values of the shortest distances of PLA spots from the nuclear surfaces between K8/K19 and K8/K19-S35E. Scale bars, 20 μm. Statistical significance was determined using one-way ANOVA followed by Dunnett’s post hoc test for (C) and an unpaired Student’s t-test for (D).

To substantiate the NMR findings, TEM imaging (Fig. 5B) was employed to visualize binding patterns and structural alterations. In both the K8/K19 + 14-3-3ζ and K8/K19-S35E + 14-3-3ζ conditions, as well as the K8/K19 + 14-3-3σ and K8/K19-S35E + 14-3-3σ conditions, “beads-on-a-string” structures with regular binding intervals were consistently observed along filaments. These observations suggest that 14-3-3ζ and 14-3-3σ binding to K8/K19 filaments is not significantly influenced by the phosphorylation at S35. Co-sedimentation assays were next performed and quantitatively confirmed the qualitative TEM findings (Fig. 5C). 14-3-3ζ binding to WT (27.75%, *P* = 0.0003) and S35E (25.50%, *P* = 0.0017) filaments was significantly greater than to the control (14.88%), corresponding to ∼1.9-fold and ∼1.7-fold increases, respectively. Similarly, 14-3-3σ showed significantly increased binding to WT (21.00%, *P* = 0.0218) and S35E (22.50%, *P*= 0.0063) filaments compared to the control (14.25%), corresponding to ∼1.5-fold and ∼1.6-fold increases. These results indicate that 14-3-3ζ and 14-3-3σ associate with K8/K19 filaments with little dependence on the S35E substitution, in contrast to the pronounced S35E-dependent reduction observed for GSK3β binding.

To determine whether the limited S35E dependence of the K19–14-3-3σ interaction observed in vitro was also observed in the cell-based assay, PLA was performed in 14-3-3σ-overexpressing 293T cells expressing either WT K19 or K19-S35E together with K8 (Fig. 5D). No significant differences were observed in the number of PLA spots per nucleus or in the mean and maximum distances of PLA spots from the nuclear surfaces between K8/K19 and K8/K19-S35E cells. Thus, no significant S35E-dependent change in the cellular proximity between K19 and 14-3-3σ was detected under these conditions.

**Fig. 6.**
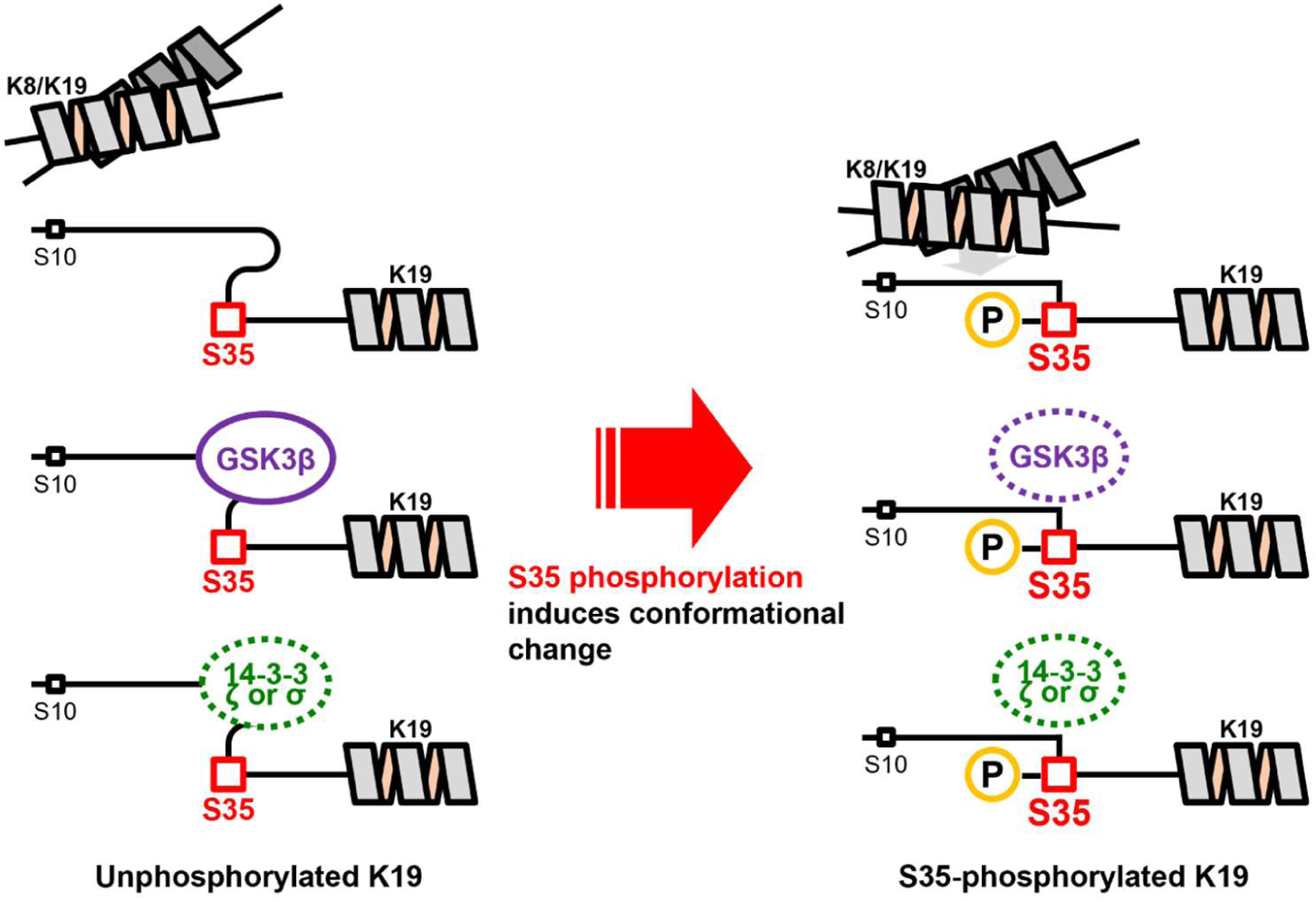
Structural and functional implications of S35 phosphorylation in the K19 head domain. The phosphomimetic S35E mutation induces pronounced conformational changes and alters interaction dynamics within the K19 head domain, expanding its functional significance beyond intracellular structural support to potential roles in cancer-associated pathways.

## Discussion

Head domains are essential for intermediate filament assembly and are the principal sites of regulatory phosphorylation. Yet they have largely resisted structural description. Here we solved the solution structure of the N-terminal portion of human K19 head domain by solution NMR and defined how phosphorylation at Ser35 reshapes it. Although keratin head domains have generally been regarded as intrinsically disordered, the K19 head domain is not uniformly so. It comprises locally structured segments with hydrophobic contacts in the derived models (H1–H3) embedded in a flexible chain. This architecture resembles the “transiently structured” behavior reported for the neurofilament light and desmin head domains, in which labile β-strand-enriched self-association within low-complexity segments governs assembly and is tuned by phosphorylation (*9*). It is also consistent with the proposal that IF head and tail domains contain low-complexity, aromatic-rich kinked segments rather than featureless random coil (*16*). Our model provides an experimental counterpart to those inferences at atomic resolution for the head domain of K19, a notably short type I keratin that is frequently overexpressed in carcinomas.

In our S35E model, the introduction of a negative charge at position Ser35 was associated with pronounced conformational rearrangement rather than a global order-to-disorder change. The domain adopted an extended, solvent-exposed configuration that repositioned the H1–H2 interface and altered the accessibility of surfaces flanking Ser35. Ser35 is the major physiological phosphorylation site in K19 (*7*), detected in cells in independent studies (*27, 36*), and reported to be targeted by Akt (*27*). The phosphorylation switch therefore offers a structural mechanism by which growth-factor signaling may be transduced into the keratin cytoskeleton, and places K19 within the paradigm in which site-specific head-domain phosphorylation, rather than bulk charge addition, dictates IF behavior (*37*).

Our result is consistent with an independent observation in cells by Ju and colleagues. They reported that Akt-mediated Ser35 phosphorylation converts K19 from an insoluble, filamentous form to a soluble, granular form (*27*). We showed, using purified keratin proteins, that a negative charge at Ser35 lowers assembly efficiency, consistent in direction with the redistribution toward the soluble form. Comparing our structural model with the recent vimentin filament structure raises the question of whether Ser35 phosphorylation occurs only before assembly, or can also take place after a filament has formed. Cryo-EM and cryo-ET of vimentin filaments showed that the disordered head domains are not displayed on the filament surface but coalesce into a fiber within the filament lumen (*12*). If type I keratin head domains are similarly sequestered, two issues follow. First, the extension of the head domain that we observed requires conformational freedom that a lumen-buried head domain may not possess, which would restrict the phosphorylation switch to the soluble, pre-assembly pool. Second, Akt would need access to Ser35 within a mature filament to modify it in situ; a buried head domain predicts that Akt acts mainly on soluble K19, not on assembled K19 in filaments. Consistent with this, K19 phosphorylation was found three decades ago to be enriched in the soluble rather than the insoluble keratin fraction (*7*). Our structural models may represent the soluble fraction of the K8/K19 before filament assembly. Comparing the phosphorylation kinetics of assembled versus soluble K8/K19 by recombinant Akt would distinguish these possibilities and determine whether head-domain phosphorylation gates filament assembly or remodels existing filaments.

Our NMR titration with K19 head peptides showed that the isolated K19 head domain binds directly to preassembled K8/K19 filaments. The head segment in our study is therefore a filament-interacting module and not merely an assembly prerequisite. Because the free head domain was added to assembled filaments, our experiment reports on binding in trans. The S35E head domain bound more avidly than wild type, as judged by the complete loss of peptide signals upon addition of K8/K19 filaments. Stronger binding of the head did not translate into better filament assembly. Co-sedimentation assays showed that the S35E mutation reduced assembly efficiency. TEM images showed that S35E-containing filaments were significantly wider than wild type, with regions of partial unraveling and irregular thickening along their length. In cells, K19-S35E mutant shortened average filament length and reduced length per junction, while apparent line width was unchanged. The unchanged line width does not conflict with the TEM measurement, because ridge-based analysis of diffraction-limited fluorescence reports the apparent width of keratin bundles, whereas TEM resolves individual filaments. One possible explanation for these observations is that a negative charge at Ser35 increases the avidity of the head domain for filament surfaces while degrading the fidelity of its association with filaments. An extended, solvent-exposed head domain presents more surface for association but is less able to make the specific head–rod contacts that canonical assembly requires. Filaments formed under these conditions would be expected to show the irregular thickening and local unraveling we observe, and fewer subunits would be recovered in the assembled fraction. A previous report that the S35A mutant also produces collapsed or shorter keratin networks (*7*) indicates that the native serine is required for normal network architecture, and that both loss of the hydroxyl and addition of negative charge at this position are disruptive. These effects are consistent with regulation of the keratin cycle, in which the balance between soluble subunits and assembled polymer is continuously adjusted (*38*).

The unphosphorylated K19 head domain bound GSK3β strongly, as judged by more pronounced peak changes in NMR spectra than those of the phosphomimetic head domain. The S35E mutation weakened GSK3β binding in vitro, reduced K19–GSK3β proximity in cells by PLA, and diminished the amount of GSK3β retained by K8/K19 filaments in co-sedimentation assay. Three independent methods converge on the same conclusion, which we regard as the central finding of this study. These results resolve an ambiguity left open by previous work. Sharma and colleagues showed that K19 binds GSK3β, sustains its inhibitory Ser9 phosphorylation of GSK3β, prevents nuclear accumulation of the active kinase, and thereby maintains cyclin D3 levels and proliferation; both Ser10 and Ser35 of K19 were required for GSK3β binding, and the S35A mutant failed to maintain cyclin D3 (*5*). In the same study, the Akt inhibitor LY294002 increased the K19– GSK3β interaction, which led the authors to propose that Ser35 might act independently of its phosphorylation, or through a kinase other than Akt. Our data support a simpler reading. Lowering Akt activity, and lowering Ser35 phosphorylation of K19, strengthens the K19–GSK3β interaction. Both observations point the same way: modification of Ser35 disengages GSK3β. One possible explanation for this apparent discrepancy is suggested by our structural models. The Ser35 side chain contributes directly to the interface, so alanine substitution can abolish a contact site. Also, adding negative charge at S35 does not simply delete that contact but remodels the surrounding region into an extended configuration. Our structural model separates these two outcomes, which mutational analysis alone cannot.

Our in vitro result extends the published mapping in one respect. Sharma and colleagues found that the rod domain was required for K19 to co-immunoprecipitate GSK3β and that the head domain alone was not sufficient in cells, yet we observe direct binding of the isolated head domain to GSK3β with purified components. The two findings are reconciled if the rod domain contributes avidity through filament assembly and local concentration, while the head domain provides the phosphorylation-sensitive contact. On this view, the interaction is bipartite, and Ser35 governs its regulated component rather than the bulk of the binding energy. In K19-positive breast cancer cells, K19 restrains GSK3β and thereby maintains cyclin D3, a D-type cyclin that active GSK3β otherwise targets for degradation (*39*), and cyclin D3 levels track with sensitivity to CDK4/6 inhibitors (*5*, *40*). A modification that releases GSK3β from the scaffold would be expected to act on this axis, although testing that prediction lies beyond the present study. A comparable arrangement has been described in another tumor type. In hepatocellular carcinoma, cytoplasmic K19 retains β-catenin and thereby influences its nuclear translocation and downstream Notch1 signaling, and Ser35 phosphorylation — detected directly in those cells — accompanies relocation of K19 to the plasma membrane (*36*). The partner differs, but the architecture is the same: cytoplasmic K19 holds a signaling protein, and modification at Ser35 is associated with release. Our structural data can provide a physical mechanism for one instance of this arrangement.

14-3-3ζ and 14-3-3σ also bound the K19 head domain, although the observed association showed little detectable dependence on the S35E substitution. NMR spectra were comparable between wild type and S35E, and PLA detected no significant S35E-dependent change in K19–14-3-3σ proximity. The conformational extension we characterized is therefore not a general switch for all partners. This is consistent with how 14-3-3 proteins engage IF proteins (*41*). For K18, phosphorylation of Ser33 is necessary but not sufficient for 14-3-3 binding, which additionally requires events coupled to mitosis (*42*). For K17, two consensus sites in the head domain, Thr9 and Ser44, each contribute: alanine substitution at either position reduces 14-3-3 binding to a similar extent, and the double mutant fails to restore cytoplasmic 14-3-3, Akt/mTOR activity or cell growth (*43*). Multivalent, combinatorial control rather than single-site switching appears to be the rule. Notably, K18 and K19 are co-expressed in the same simple epithelia yet differ in their basal phosphorylation and in their sensitivity to phosphatase inhibition (*7*), indicating that type I keratins are regulated individually rather than as a class. These head-domain sites, in K17 as in K19, have been mapped by alanine substitution. That approach establishes that a serine or a threonine residue is required, but does not distinguish a side-chain contact from a phosphorylation-dependent one. The structural analysis reported here was designed to address that distinction more precisely. Ser10 and Ser22 lie near Ser35 in sequence and cluster with it in our three-dimensional model. Since a 14-3-3 dimer engages two phosphosites, a high-affinity K19–14-3-3 interface may require co-modification of Ser35 with one or more neighbors. Testing this will require head domains bearing genuine phosphoserine at defined combinations of these positions, which lies beyond the scope of the present study.

Several limitations bound the interpretation of our results. First, the S35E mutation is a phosphomimetic, not a phosphoserine. Glutamate carries a single negative charge and a planar carboxylate, whereas phosphoserine carries up to two negative charges in tetrahedral geometry. This distinction may matter for actual binding with GSK3β, whose phosphate-recognition pocket is tailored to a genuine phosphate. Our results are most rigorously read as showing that a negative charge at Ser35 restructures the head domain and weakens GSK3β engagement. Second, our NMR work used the isolated head domain with short peptides. This makes residue-level analysis tractable but removes the rod and tail domains and the assembled-filament context. Therefore, the extended conformation with S35E is established for the free head domain.

Third, the cellular experiments used 293T cells, which provide a keratin-poor background suited to reconstituting a defined K8/K19 pair but do not reproduce the signaling environment of a breast carcinoma cell. Fourth, the increased filament width seen by TEM is compatible with either altered subunit packing within single filaments or enhanced filament–filament association. The regions of partial unraveling in the same images favor altered packing, but our data do not exclude a contribution from lateral association. Fifth, we did not measure nuclear GSK3β, cyclin D3 levels or proliferation in the S35E-expressing 293T cells. The connection between the structural switch described here and the cell-cycle consequences reported in the literature therefore remains inferential.

Several experiments would extend this work. One approach would be to generate a pSer35 head domain, either synthetically or enzymatically with a kinase such as Akt, and compare it with S35A as a matched control. This would test whether authentic phosphorylation reproduces both the conformational extension and the loss of GSK3β binding, and would separate the effect of negative charge from that of losing the side-chain hydroxyl. In addition, measuring nuclear GSK3β, cyclin D3 stability and CDK4/6-inhibitor sensitivity in parallel wild-type, S35A and S35E lines would establish whether the structural switch carries functional consequences. Pharmacological manipulation of Akt in the same lines would test pathway dependence.

Determining whether Akt can phosphorylate Ser35 within assembled K8/K19 filaments, or only in the soluble pool, would resolve the accessibility question raised above. Cryo-ET of K8/K19 filaments, following the approach applied to vimentin (*12*), would show directly whether type I keratin head domains are lumen-sequestered. Finally, the K19 head domain shares sequence features with other type I keratins while differing at residues that may set partner specificity. Systematic comparison of head domains across the keratin family offers a route to understanding how a shared architectural module achieves partner-selective signaling.

In conclusion, our study provides a structural model of the N-terminal head domain of human K19 using solution NMR spectroscopy and explores its structural dynamics. Our results suggest that site-specific phosphorylation at Ser35 plays a role in modulating K19 interactions with cancer-related signaling proteins, such as GSK3β. While the precise mechanism remains elusive, phosphorylation-induced structural alterations in the head domain of K19 may influence interactions between K19 and its binding partners, suggesting a possible mechanism for signaling-related regulation of cellular processes beyond their established role as a structural scaffold.

## Materials and Methods

### Peptide Synthesis and Purity Verification

All the peptides used for NMR spectroscopy in this study were synthesized by Peptron Inc. (Daejeon, Republic of Korea) using solid-phase peptide synthesis (SPPS) based on the sequence of Keratin, type I cytoskeletal 19 (P08727 · K1C19_HUMAN). The identity and purity of each peptide were confirmed by analytical high-performance liquid chromatography (HPLC) and matrix-assisted laser desorption/ionization time-of-flight mass spectrometry (MALDI-TOF MS). All peptides were obtained at >90% purity. NMR spectral analysis further supported the identities of the synthesized peptides.

### NMR Experiments and Structural Analysis

All NMR data were collected using an Avance III HD 850 MHz spectrometer equipped with a cryogenic HCN probe (Bruker). For all experiments, the acquisition temperature was maintained at 298 K, and the sample volume was set to 300 μL, using a 5-mm Shigemi tube (Sigma-Aldrich). Peptides for solution structural modeling were prepared at a concentration of 3.3 mg/mL in a buffer containing 20 mM sodium phosphate (pH 7.4) and 7% deuterium oxide (D2O). The following experiments were performed for signal assignment and structural calculation: 1D ^1^H, 2D ^1^H-^13^C HSQC, 2D ^1^H-^1^H TOCSY, and 2D ^1^H-^1^H ROESY. To investigate the interaction of H1H2H3 and H1H2H3-S35E with K8/K19 filaments, GSK3β, 14-3-3 proteins, and GST, peptide samples were prepared at a concentration of 100 μM in a buffer containing 5 mM sodium phosphate (pH 7.4) and 7% D2O. This lower concentration was necessary to maintain at least a 1:1 molar ratio between the peptides and interacting partners, though the availability of some protein candidates (e.g., K8/K19 filaments and GSK3β) was limited. After acquiring initial 1D ^1^H and 2D ^1^H-^13^C HSQC spectra of the peptide-only sample, an equivalent amount of the interacting partner was added. Subsequently, 1D ^1^H and 2D ^1^H-^13^C HSQC spectra were re-recorded. To distinguish peptide resonances from those of the interacting proteins, these same spectra were also acquired for the protein-only samples.

The NMR data were acquired and processed using TopSpin 3.2 software (Bruker). Chemical shift assignments and subsequent analysis were performed with POKY software (*44*). Each peak in the NMR spectra was manually identified. Chemical shift perturbations (CSPs) were calculated using a predefined equation.

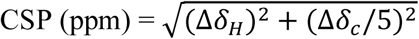

The chemical shift differences were analyzed and visualized with a custom Python program utilizing the pandas, numpy, seaborn, re, and matplotlib libraries.

Distance constraints for structural determination were derived from peaks observed in 2D ^1^H-^1^H ROESY spectra. Angle constraints were extracted from chemical shift data, including Cα, Cβ, CO, HN, N, Hα, and Hβ shifts, using TALOS-N software (*45*). The initial peptide structure model was generated using Xplor-NIH in combination with POKY (*46*). Refinement of the structure was performed using Xplor-NIH, TALOS-N, and Cyana (*47*). PyMOL was employed for structure visualization and analysis (*48*). The contact map was generated based on distances between the Cβ atoms of paired residues in the average structure model, with a cutoff value of 7.5 Å. For residues where Cβ data were unavailable (e.g., glycines), Cα was used as an alternative. The contact map was computed and visualized with a custom Python program, utilizing pandas, numpy, seaborn, and matplotlib Python 3 libraries.

### Transformation and protein expression

Recombinant plasmids encoding the target proteins were introduced into *Escherichia coli* Rosetta (DE3 pLysS) competent cells using a standard heat-shock method. The following plasmids were used for expression: pET29 for Keratin 8, pET3d for Keratin 19, pGEX for GSK3β, pEXP for GST, and pET28b for both 14-3-3ζ and 14-3-3σ. After transformation, cells were spread onto LB agar plates containing the appropriate selection antibiotics. Kanamycin was used for pET28b and pET29 vectors, whereas ampicillin was used for pET3d, pGEX, and pEXP constructs. Plates were incubated overnight at 37°C. Individual colonies were picked and cultured in LB medium supplemented with the corresponding antibiotic to establish seed cultures. Once the OD600 of these cultures reached 0.6–0.8, protein expression was induced by the addition of 1 mM IPTG. Depending on the solubility of the expressed proteins, cultures were incubated either at 16°C to promote soluble protein expression or at 37°C for proteins prone to form inclusion bodies. After 16 to 20 hours of shaking incubation, bacterial cells were harvested by centrifugation at 4,000 × g for 30 minutes at 4°C. The pellets were then stored at -80°C until further use.

### Purification of Keratin Proteins

Due to their inherent insolubility, keratin proteins predominantly accumulate as inclusion bodies during overexpression. To isolate them, transformed *E. coli* cells were first collected and resuspended in lysis buffer comprising 50 mM Tris-HCl (pH 8.0), 1 mM EDTA, 2 mM DTT, and 1 mM PMSF. Cell disruption was achieved through sonication, after which the lysate was subjected to centrifugation at 17,200 × g for 30 minutes at 4°C to separate soluble and insoluble fractions. The supernatant was discarded, and the resulting pellet, enriched in inclusion bodies, underwent a series of washing steps to eliminate cellular contaminants. The initial wash utilized inclusion body wash buffer I (50 mM Tris-HCl (pH 8.0), 0.2 M NaCl, 1 mM EDTA, 2 mM DTT, 1 mM PMSF, and 1% NP-40), followed by centrifugation under identical conditions. The pellet was then treated with inclusion body wash buffer II (50 mM Tris-HCl (pH 8.0), 0.5 M NaCl, 1 mM EDTA, 2 mM DTT, 1 mM PMSF, 0.5% Triton X-100, and 0.5% Tween-20), and centrifuged again to yield a washed inclusion body pellet (tight pellet). This pellet was solubilized in denaturing buffer A (6 M urea, 50 mM Tris-HCl (pH 8.0), 1 mM EDTA, 2 mM DTT, 1 mM PMSF) and passed through a filter to remove any remaining insoluble material. The solubilized keratin solution was subjected to ion-exchange chromatography using an ÄKTA start™ system equipped with a HiTrap Q column (Cytiva). A linear gradient elution was performed using buffer B (6 M urea, 50 mM Tris-HCl (pH 8.0), 1 mM EDTA, 2 mM DTT, 1 mM PMSF, 300 mM guanidinium chloride). Protein-containing fractions were monitored via SDS-PAGE and pooled based on purity. Quantification of the purified keratin was carried out using the Bradford protein assay. For filament assembly, type I and type II keratin isoforms were combined in equimolar amounts and incubated overnight at 4°C. Following assembly, the keratin filaments were further purified by reapplying the mixture to a HiTrap Q column to enrich for fully assembled filamentous complexes.

### Filament Assembly of Keratin Proteins

Purified keratin proteins were assembled into filaments through a stepwise dialysis procedure designed to promote gradual refolding and reconstruction of their native structure. SDS-PAGE-confirmed protein fractions were pooled and loaded into cellulose dialysis membranes. The initial dialysis was carried out at room temperature for 4 hours against a buffer containing 9 M urea, 25 mM Tris-HCl (pH 7.4), and 25 mM β-mercaptoethanol. This was followed by a second dialysis step at 4°C for 6 hours using a buffer containing 2 M urea, 5 mM Tris-HCl (pH 7.4), and 5 mM β-mercaptoethanol to further facilitate protein refolding. In the final step, the sample was dialyzed against 5 mM sodium phosphate (pH 7.4) containing 5 mM β-mercaptoethanol at 4°C for 6 hours, followed by overnight incubation to ensure complete urea elimination.

### Purification of GSK3β and GST

Recombinant GSK3β and GST proteins were expressed from *E. coli* strains harboring the pGEX-GSK3β and pEXP-GST plasmids, respectively. Following cell harvest, pellets were resuspended in phosphate-buffered saline (PBS) supplemented with 1 mM EDTA, 2 mM DTT, and 1 mM PMSF. Cells were lysed via sonication, and the resulting lysate was clarified by filtration to remove insoluble debris. The clarified lysate was applied to a Pierce™ Glutathione Agarose column pre-equilibrated with the same PBS-based buffer. After protein binding, the resin was washed thoroughly with washing buffer (PBS containing 1 mM EDTA, 2 mM DTT, and 1 mM PMSF) to eliminate nonspecifically bound proteins. Bound GST-tagged proteins were eluted using a buffer containing 50 mM Tris-HCl (pH 8.0) and 10 mM reduced glutathione. Fractions containing the protein of interest were identified by SDS-PAGE and pooled accordingly. The GSK3β-containing eluate was concentrated using a centrifugal filter device, followed by an additional purification step via size-exclusion chromatography (SEC) using an ÄKTA pure™ system with a Superose™ 6 Increase 10/300 GL column (Cytiva). SEC fractions were analyzed by SDS-PAGE to confirm purity, and those containing highly purified GSK3β were combined and concentrated once more. During the final concentration step, the buffer was exchanged to 5 mM sodium phosphate (pH 7.4) containing 5 mM β-mercaptoethanol to suit downstream applications.

### LC-MS/MS Analysis and Western Blot of Purified GSK3β

LC-MS/MS analysis was performed by eMass Co. (Korea). Proteins were subjected to in-gel digestion using trypsin or chymotrypsin after standard reduction, alkylation, and dehydration steps. Peptides were extracted, desalted using Sep-Pak C18 cartridges, dried by SpeedVac, and reconstituted in 0.1% formic acid before injection. LC separation was carried out using a Thermo Dionex Ultimate 3000 UHPLC system with an ACQUITY UPLC BEH 130 C18 column (1.7-μm particle size, maintained at 50 °C). MS analysis was conducted on an AB Sciex TripleTOF 5600+ using nitrogen gas, with a mass range of 50–2500 m/z. Data acquisition used Analyst TF 1.7.1 and data interpretation was performed using ProteinPilot 4.5 and PeakView 2.20 software with enzyme-specific parameters; proteins were identified based on >95% confidence in de novo or reproducible peptide matching.

Western blotting was performed to confirm the presence of GSK3β in the purified sample. Proteins were separated by SDS-PAGE and transferred onto a PVDF membrane (Amersham Protran, Cytiva, Marlborough, MA, USA). The membrane was probed with a primary antibody against GSK3β (ABclonal, rabbit polyclonal antibody, A2081), followed by incubation with an HRP-conjugated goat anti-rabbit immunoglobulin G secondary antibody (AP132P, EMD Millipore). Detection was carried out using a chemiluminescence imaging system (WSE-6200 LuminoGraph II, ATTO Korea).

### Purification of 14-3-3

Recombinant 14-3-3ζ and 14-3-3σ proteins were expressed using *E. coli* strains transformed with pET28b vectors encoding each isoform. Following bacterial harvest, cell pellets were resuspended in lysis buffer (50 mM Tris-HCl, pH 8.0, 0.5 M NaCl, 10 mM imidazole, 1 mM PMSF) and disrupted via sonication. The resulting lysate was filtered to remove particulate matter, and the cleared supernatant was subjected to affinity purification. Nickel-affinity chromatography was conducted on an ÄKTA pure™ system (Cytiva) using a HisTrap column pre-equilibrated with the same lysis buffer. After sample loading, the column was washed with equilibration buffer to eliminate nonspecifically bound proteins. Elution was achieved through a stepwise gradient of buffer containing increasing concentrations of imidazole (elution buffer: 50 mM Tris-HCl, pH 8.0, 0.5 M NaCl, 0.5 M imidazole, 1 mM PMSF). Protein-containing fractions were analyzed by SDS-PAGE to evaluate expression and purity. Fractions enriched in 14-3-3ζ or 14-3-3σ were pooled and concentrated using a centrifugal filtration unit. Simultaneously, buffer exchange was performed into 5 mM sodium phosphate (pH 7.4) supplemented with 5 mM β-mercaptoethanol.

### Transmission Electron Microscopy (TEM)

To assess keratin filament assembly and characterize filament ultrastructure, and visualize interactions with associated proteins such as GSK3β and 14-3-3 isoforms, negative-stain transmission electron microscopy was employed (Fig. S3). Keratin filaments were diluted to a final concentration of 0.05 mg/mL, and 10 μL of each sample was deposited onto Formvar/carbon-coated copper grids (300 mesh, Ted Pella, 01753-F). For protein-binding assays, keratin filaments were incubated with GSK3β or 14-3-3 proteins at a 1:10 molar ratio (binding partner: keratin), prior to staining. Samples were stained with 1% uranyl acetate to enhance image contrast. TEM imaging was carried out using an FEI Tecnai G2 electron microscope (operated at 120 kV) housed at the Korea Brain Research Institute (KBRI). Images were captured at multiple magnifications (7,800×, 14,500×, 29,000×, and 50,000×) to examine filament morphology and protein binding. DigitalMicrograph software (Gatan) was used for image acquisition and subsequent analysis.

For quantification of filament width, images acquired at 29,000× magnification were selected. Four TEM fields were randomly chosen per sample, and filament diameters were measured perpendicular to the longitudinal axis using a calibrated line measurement tool. A total of 100 individual measurements were performed for each experimental condition. Due to unequal variances between groups (F = 9.728, *P* < 0.0001), Welch’s t-test was used to compare filament widths of K8/K19 WT and S35E phosphomimetic mutants. The average width of WT filaments was 7.553 nm, while S35E filaments exhibited an increased mean width of 10.73 nm. The mean difference between the groups was 3.182 ± 0.2661 nm (95% CI: 2.655–3.709; *P* < 0.0001, two-tailed), corresponding to a large effect size (η² = 0.5454), indicative of a significant structural change associated with the S35E mutation.

### Sedimentation Assay of Keratin Filaments and K19-Binding Partners

To evaluate the efficiency of keratin filament assembly and their interactions with potential binding partners, sedimentation assays were performed under ultracentrifugation. Reconstituted keratin and keratin-binding partner mixtures (120 μL) at a final keratin concentration of 0.25 mg/mL were centrifuged using an Airfuge system at 70,000 rpm (approximately 96,680 × g) for 1 hour at room temperature. Following centrifugation, supernatant and pellet fractions were carefully separated. The pellet was resuspended in SDS sample buffer (Biosesang), and both fractions were analyzed by SDS-PAGE to confirm filament assembly and sedimentation behavior.

Band intensities on SDS-PAGE gels were quantified using ImageJ software. For each binding partner condition, the intensities of both pellet and supernatant bands were measured. To account for variability between replicates, all intensity values were normalized to the total signal (pellet + supernatant), which was defined as 100%. Relative binding was expressed as the ratio of pellet intensity to total protein. Statistical analysis was conducted using Welch’s t-test to compare filament assembly efficiency, and one-way ANOVA followed by Dunnett’s post-hoc test for multiple comparisons to evaluate relative binding levels. All statistical analyses were conducted using GraphPad Prism version 11.1.0 (GraphPad Software, San Diego, CA, USA).

### Plasmid Construction

For expression in 293T cells, the coding sequences of human K8 (WT) and human K19 (WT, S35E, and S35A) were subcloned into pcDNA3.1. Each coding sequence was amplified by PCR using Pfu DNA polymerase (Bioneer) from the corresponding bacterial expression plasmids (pET29-K8 WT and pET3d-K19 WT, S35E, and S35A). Since the S35E and S35A substitutions were already present in the parental pET3d templates, the mutant coding sequences were transferred to pcDNA3.1 without additional site-directed mutagenesis. The forward primers were designed to append a Kozak consensus sequence and an additional glycine codon immediately downstream of the initiating ATG codon. The pcDNA3.1 vector was linearized with NheI and XhoI, and each amplicon was joined to the linearized vector using In-Fusion® Snap Assembly Master Mix (Takara Bio) according to the manufacturer’s instructions. The assembled products were transformed into E. coli DH5α using a standard heat-shock method, and transformants were selected on LB agar containing ampicillin. Plates were incubated overnight at 37°C, and individual colonies were screened by colony PCR. Positive colonies were inoculated into LB broth containing ampicillin and cultured overnight at 37°C with shaking. Plasmid DNA was isolated from the cultures and verified by Sanger sequencing.

### Cell Culture and Generation of Stable Cell Lines

293T cells were maintained in high-glucose DMEM (Gibco, Waltham, MA, USA; 11965092) supplemented with 10% fetal bovine serum (Corning, Corning, NY, USA; 35-015-CV) and 1% penicillin/streptomycin at 37 °C under 5% CO₂. To generate stable cell lines, cells were transfected with pcDNA3.1-based plasmids encoding 14-3-3σ or GSK3β, or with the empty pcDNA3.1 vector as a control, using Lipofectamine 3000 (Invitrogen, Carlsbad, CA, USA; L3000001) per the manufacturer’s instructions. At 48 h post-transfection, cells were selected in medium containing 800 µg/mL G418 (Geneticin; InvivoGen, San Diego, CA, USA; ant-gn-1). After 2 weeks, single clones were randomly isolated, expanded, and maintained under continuous G418 selection, and stable transgene expression was confirmed by western blotting (Fig. S5) before downstream experiments.

### Transfection and Immunofluorescence

Stable 293T cells were seeded onto sterile glass coverslips and co-transfected with wild-type K8 together with either wild-type K19 or the K19 S35E mutant at a K8:K19 ratio of 3:1, using Lipofectamine Plus reagent (Invitrogen, Carlsbad, CA, USA; 15338100). At 48 h post-transfection, cells were fixed in 100% cold methanol and blocked with 2% normal goat serum in PBS. Samples were incubated overnight at 4 °C with a recombinant rabbit monoclonal anti-cytokeratin 8 antibody (Invitrogen, Waltham, MA, USA; MA5-32118) and a mouse monoclonal anti-cytokeratin 19 antibody (Santa Cruz Biotechnology, Dallas, TX, USA; sc-6278), then with FSD 594-conjugated goat anti-rabbit IgG (BioActs, Incheon, South Korea; RSA1295) and FSD 488-conjugated goat anti-mouse IgG (BioActs, Incheon, South Korea; RSA1145). Nuclei were counterstained with DAPI, and coverslips were imaged on a Carl Zeiss LSM 900 confocal microscope (Carl Zeiss, Oberkochen, Germany). Keratin filament organization was quantified by ridge detection using Fiji (*49*, *50*).

### Proximity Ligation Assay (PLA)

In situ protein proximity was detected using the Duolink In Situ Red Starter Kit Mouse/Rabbit (Sigma-Aldrich, St. Louis, MO, USA; DUO92101) according to the manufacturer’s instructions (*51*). To visualize the interactions between 14-3-3σ and K19 and between GSK3β and K19, cells were fixed, permeabilized, and blocked, then incubated with the corresponding pairs of mouse and rabbit primary antibodies, followed by anti-mouse PLUS and anti-rabbit MINUS PLA probes, ligation, and rolling-circle amplification. Nuclei were counterstained, and coverslips were mounted with the supplied medium. PLA signals were imaged on a Carl Zeiss LSM 900 confocal microscope, and the number and spatial distribution of PLA spots were quantified using IMARIS (Bitplane, Zurich, Switzerland). Three-dimensional reconstruction of the PLA signals was also performed using IMARIS (Fig. S6).

### Image Analysis and Statistics for Cell-Based Assays

Statistical analyses for the cell-based assays were performed using GraphPad Prism. Comparisons between two independent groups were assessed by unpaired Student’s t-test. Keratin filament morphology data are presented as violin plots showing minimum and maximum values, whereas PLA data (spot counts and distances) are presented as dot plots of individual data points overlaid with the mean ± SD. *P* < 0.05 was considered statistically significant.

## Funding

This work was supported by the National Research Foundation of Korea (NRF) (No. RS-2026-25474533 to Lee CH and 2020R1I1A2074335 to Kim JH). PAC was supported by grant R01 AR079418 from the National Institutes of Health in U.S.A. This work was also supported by the InnoCORE program of the Ministry of Science and ICT (26-InnoCORE-07). NMR experiments were supported by the Institute for Basic Science of Korea (No. IBS-R034-D1 to Ko YH) funded by the Korean government (MSIT). TEM data were acquired at Brain Research Core Facilities in KBRI. We thank Dr. Ji Young Mun for her precious feedback for TEM experiments. We also thank Professor Young-Sam Lee and Dr. Jinju Jeong for their valuable feedback on this project. Software licenses for PyMOL and GraphPad Prism were funded by DGIST Supercomputing AI Education and Research Center.

## Author contributions

Ji Y and Jeong M conducted NMR analyses and structural modeling in this study. Kim J and Park C carried out the experiments involving proteins, including the purification and assembly of K8/K19 keratin filaments, GSK3β, and 14-3-3. Yeom J carried out the studies involving keratin filaments. Han MA conducted supporting experiments for GSK3β. Lee SH contributed to imaging the proteins using TEM. Hong E contributed to the protein biochemistry experiments. Kim JH designed structural experiments and conducted NMR spectroscopy. Ko YH contributed to setting up NMR experiments. Coulombe PA contributed to designing the 14-3-3 studies and assisted with biological interpretation. Lee CH launched the project, designed the experimental scheme, and conducted structural analyses for this study. All authors participated in data analyses, manuscript writing, and revisions.

## Competing interests

The authors declare that they have no competing interests.

## Data and materials availability

The datasets related to the structures and NMR in this study are available in the RCSB PDB (https://www.rcsb.org/) with the accession codes listed in Table S1. The data supporting the findings of this study are available from the corresponding authors, Lee CH or Kim JH, upon reasonable request.

## Notes

### Competing Interest Statement

The authors have declared no competing interest.

