## Supplementary Materials for "Phosphorylation-dependent conformational change in the Keratin 19 head domain: Implications for filament assembly and binding to GSK3β and 14-3-3"

Youngheum Ji *et al.*

**This PDF file includes:**

Tables S1 to S3
Figures S1 to S6

### PDB Deposits and Structural Statistics

**Table S1. Summary of deposited PDB structures.**

| PDB ID | Label | Structural information |
|---|---|---|
| 8ZG2 | H1 | Human Keratin 19 head domain segment Y6-G16 in solution |
| 8ZG3 | H2 | Human Keratin 19 head domain segment L18-G28 in solution |
| 8Y3S | H3 | Human Keratin 19 head domain segment G28-G38 in solution |
| 8ZUG | H1H2 | Human Keratin 19 head domain segment Y6-G28 in solution |
| 8XTO | H2H3 | Human Keratin 19 head domain segment L18-G38 in solution |
| 8XTN | H1H2H3 | Human Keratin 19 head domain segment Y6-G38 in solution |
| 8ZG4 | H1-S10E | Human Keratin 19 head domain S10E phosphomimetic mutant segment Y6-G16 in solution |
| 8ZP3 | H1H2H3-S35E | Human Keratin 19 head domain S35E phosphomimetic mutant segment Y6-G38 in solution |

Structural calculations were performed using solution NMR data for three short peptides (H1, H2, H3), two longer peptides (H1H2, H2H3), and the full-length construct H1H2H3. Additionally, two phosphomimetic mutants, H1-S10E and H1H2H3-S35E, were analyzed to explore the impact of serine phosphorylation on the head domain of keratin 19 (Table S1).

**Table S2. Structural ensemble statistics determined in this study.**

| | | H1 | H2 | H3 | H1H2 | H2H3 | H1H2H3 | H1-S10E | H1H2H3 -S35E |
|---|---|---|---|---|---|---|---|---|---|
| Intra residue NOE | | 123 | 138 | 93 | 205 | 174 | 238 | 172 | 214 |
| Inter residue NOE | | 73 | 83 | 109 | 109 | 198 | 251 | 139 | 134 |
| Total NOE restraints | | 196 | 221 | 202 | 314 | 372 | 489 | 311 | 348 |
| Total experimental restraints | | 70 | 72 | 84 | 132 | 121 | 180 | 107 | 111 |
| Total backbone dihedral angle restraints | | 10 | 6 | 5 | 16 | 17 | 33 | 8 | 33 |
| Average number of | Distance restraint violations (>0.5 Å) | 0 | 0 | 0 | 0 | 0 | 0 | 0.1 | 0 |
| | Dihedral angle violations (>10.0°) | 0 | 0 | 0 | 0 | 0 | 0 | 0 | 0 |
| Backbone RMSD (ordered residue) | | 0.07 | - | 0.3 | - | 1.2 | 0.8 | 0.81 | 0.36 |
| Ordered residue | | 7-14 | - | 28-38 | - | 22-38 | 6-37 | 6-14 | 28-36 |
| All heavy atom RMSD (full length) (Å) | | 0.7 | 0.8 | 1.5 | 0.9 | 2.5 | 2.9 | 1.4 | 1.7 |
| Rama (%) | Favoured | 88 | 97 | 84 | 92 | 93 | 91 | 61 | 94 |
| | Allowed | 12 | 3 | 8 | 6 | 4 | 8 | 35 | 6 |
| | Outliers | 0 | 0 | 8 | 2 | 3 | 1 | 4 | 1 |
| Procheck raw score (φ and ψ/all dihedral angle) | | -0.52 /0.00 | -0.16 /-0.02 | -1.02 /-1.00 | -0.04 /0.08 | -0.67 /-0.60 | -0.58 /-0.43 | -1.35 /-0.41 | -0.58 /-0.43 |
| Procheck Z-score (φ and ψ/all dihedral angle) | | -1.73 /0.00 | -0.31 /-0.12 | -3.70 /-5.91 | 0.16 /0.47 | -2.32 /-3.55 | -1.97 /-2.54 | -5.00 /-2.42 | -1.97 /-2.54 |

This table presents a summary of the NMR-based statistical analyses and structural parameters for the eight deposited PDBs from this study (Table S2). For each structure, the 20 lowest-energy models were selected for ensemble construction and model deposition. All models underwent numerous iterative refinement cycles during an optimization process to ensure accuracy. Despite the flexible nature of these structures, the models for each structure converged to a consistent state. The structural validity of the calculated ensembles was assessed by evaluating the number of distance restraints, dihedral angle restraints, RMSD, and Ramachandran plot (Rama) statistics, all of which indicate data consistency and model accuracy. Backbone RMSD and heavy atom RMSD were both within 3 Å, indicating high-resolution structures and strong convergence. These findings confirm that the calculated models accurately represent the anterior half of the K19 head domain at high atomic resolution.

### 14-3-3 Binding Prediction for the K19 Head Domain

**Table S3.** Sequence-based prediction of phospho-Ser/Thr binding scores to 14-3-3 in the K19 head domain (M1-N79), evaluated for serine and threonine sites.

| Site | Peptide [-6:4] | ANN | PSSM | SVM | Consensus |
|---|---|---|---|---|---|
| 2 | -----M t SYSY | 0.200 | -0.142 | -0.857 | -0.266 |
| 3 | ----MT s YSYR | 0.118 | -0.446 | -1.096 | -0.475 |
| 5 | --MTSY s YRQS | 0.275 | -0.009 | -0.401 | -0.045 |
| 9 | SYSYRQ s SATS | 0.167 | -0.028 | -0.921 | -0.261 |
| 10 | YSYRQS s ATSS | 0.198 | 0.651 | -0.591 | 0.086 |
| 12 | YRQSSA t SSFG | 0.246 | 0.276 | -0.588 | -0.022 |
| 13 | RQSSAT s SFGG | 0.218 | -0.093 | -0.599 | -0.158 |
| 14 | QSSATS s FGGL | 0.149 | -0.099 | -0.666 | -0.205 |
| 22 | GGLGGG s VRFG | 0.082 | -0.282 | -1.188 | -0.463 |
| **35** | **VAFRAP s IHGG** | **0.642** | **0.727** | **0.309** | **0.559** |
| 40 | PSIHGG s GGRG | 0.091 | -0.298 | -1.305 | -0.504 |
| 46 | SGGRGV s VSSA | 0.204 | 0.562 | -0.402 | 0.121 |
| 48 | GRGVSV s SARF | 0.267 | 0.215 | -0.664 | -0.061 |
| 49 | RGVSVS s ARFV | 0.099 | -0.246 | -1.571 | -0.573 |
| 54 | SSARFV s SSSS | 0.245 | 0.530 | -0.556 | 0.073 |
| 55 | SARFVS s SSSG | 0.093 | -0.092 | -1.471 | -0.490 |
| 56 | ARFVSS s SSGA | 0.139 | 0.119 | -0.849 | -0.197 |
| 57 | RFVSSS s SGAY | 0.442 | 0.241 | -0.188 | 0.165 |
| 58 | FVSSSS s GAYG | 0.129 | -0.031 | -0.921 | -0.275 |
| 70 | GYGGVL t ASDG | 0.194 | 0.017 | -1.017 | -0.269 |
| 72 | GGVLTA s DGLL | 0.334 | 0.194 | -0.578 | -0.017 |

The full amino acid sequence of the human keratin 19 head domain was input for analysis to identify the most probable phosphorylation site for 14-3-3 binding (Table S3). Three predictive algorithms, artificial neural network (ANN), position-specific scoring matrix (PSSM), and support vector machine (SVM), were applied, using the same methodology described in (*33*). Among these, only the serine 35 (S35) site exceeded the threshold scores for ANN (0.55) and SVM (0.25), while the PSSM score (0.727) closely approached its threshold of 0.80. All other serine or threonine sites scored below the thresholds across all algorithms, supporting the functional importance of S35 as the primary 14-3-3 binding site.

### NMR Structures of H1H2, H2H3, and H1-S10E

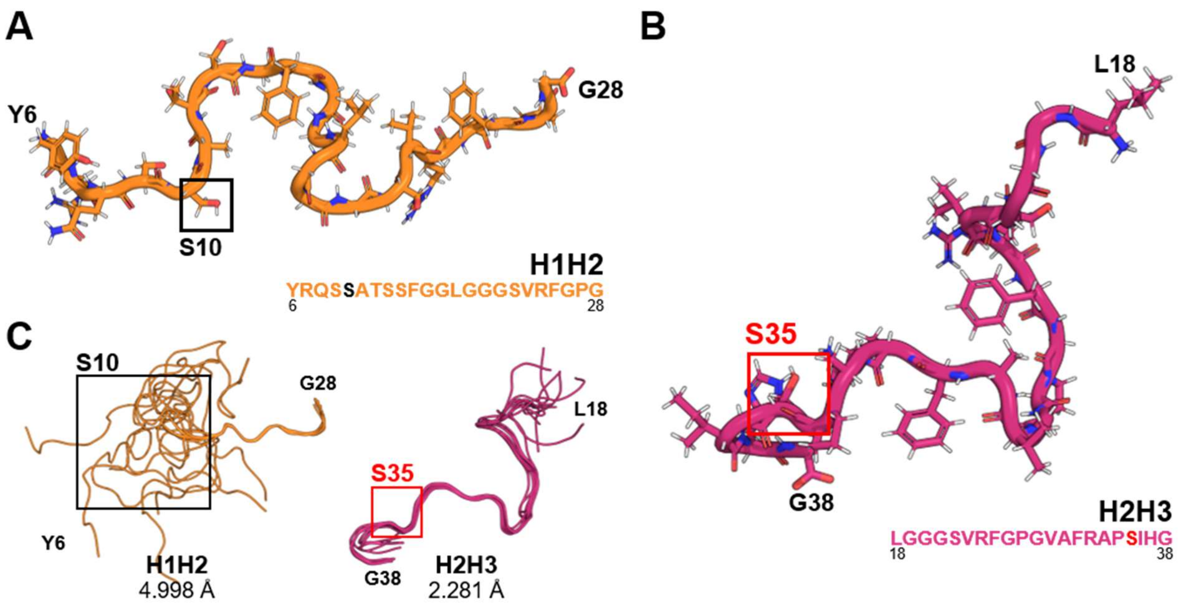


**Figure. S1. Calculated NMR models of H1H2 and H2H3. (A, B)** Representative NMR-derived structural models of H1H2 and H2H3, respectively, corresponding to the minimum-energy structures among the top 20 calculated models. **(C)** Superimposition of the 10 lowest-energy NMR-derived structural models of H1H2 and H2H3. The RMSD values were calculated over all residues.

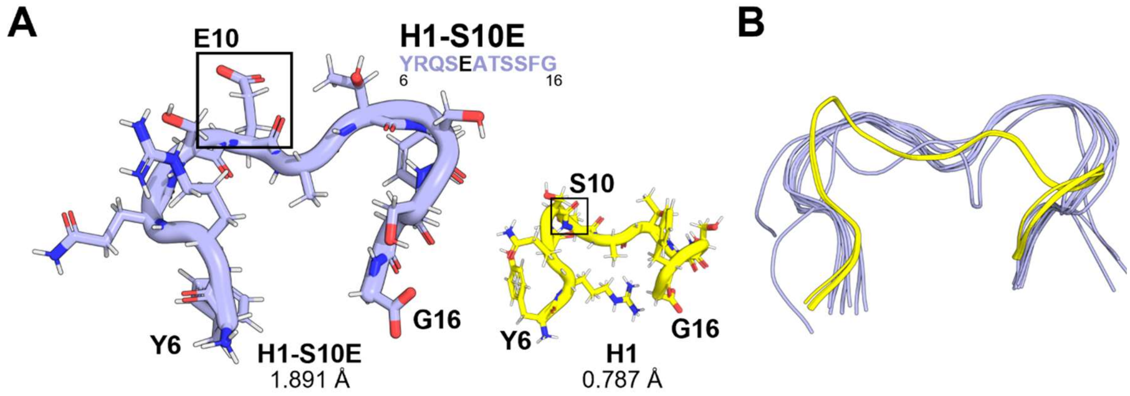


**Figure. S2. Structural comparison of the phosphomimetic mutant H1-S10E and H1. (A)** Representative NMR-derived structural models of H1-S10E and H1. The S10E substitution produced only modest structural changes in H1, in contrast to the pronounced conformational changes observed for the S35E substitution in H1H2H3. The RMSD values were calculated over all residues. **(B)** Superimposition of the 10 lowest-energy NMR-derived structural models of H1-S10E and H1. S10 did not meet the prediction thresholds for 14-3-3 binding (Table S3).

### TEM Analysis of K19-Binding Partners

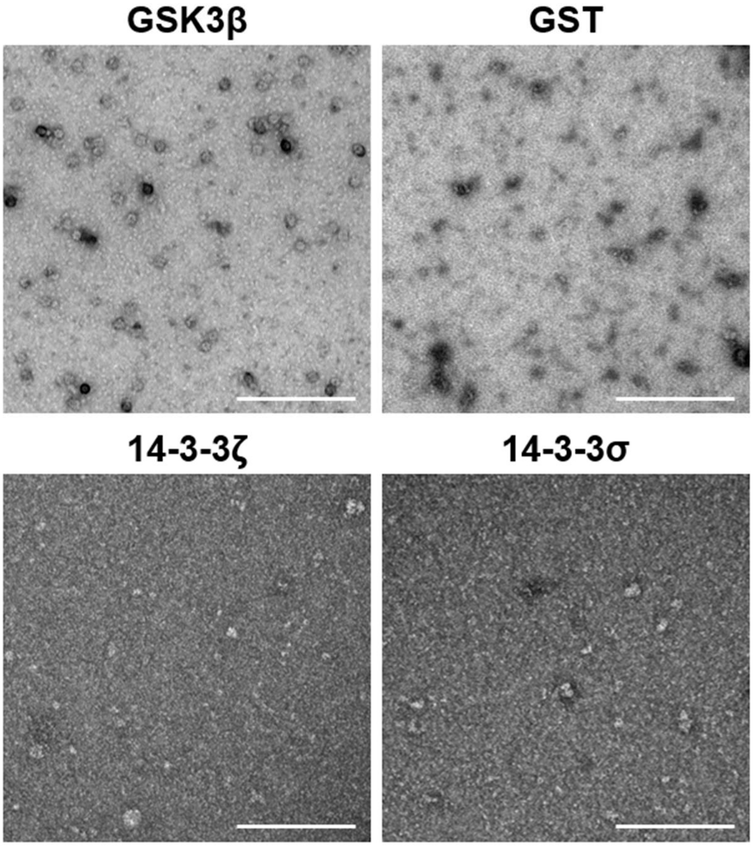


**Figure. S3. TEM Analysis of GSK3β, GST, 14-3-3ζ, and 14-3-3σ was performed to assess whether GSK3β, GST, 14-3-3ζ, and 14-3-3σ exhibit macromolecular aggregation or filamentous assembly.** All samples were imaged at 29,000× magnification. The results indicate that none of the four proteins form higher-order aggregates, such as filamentous structures or ordered assemblies, confirming their existence as individual molecules under TEM conditions. Scale bars, 200 nm.

### Quality Assessment of Purified GSK3β

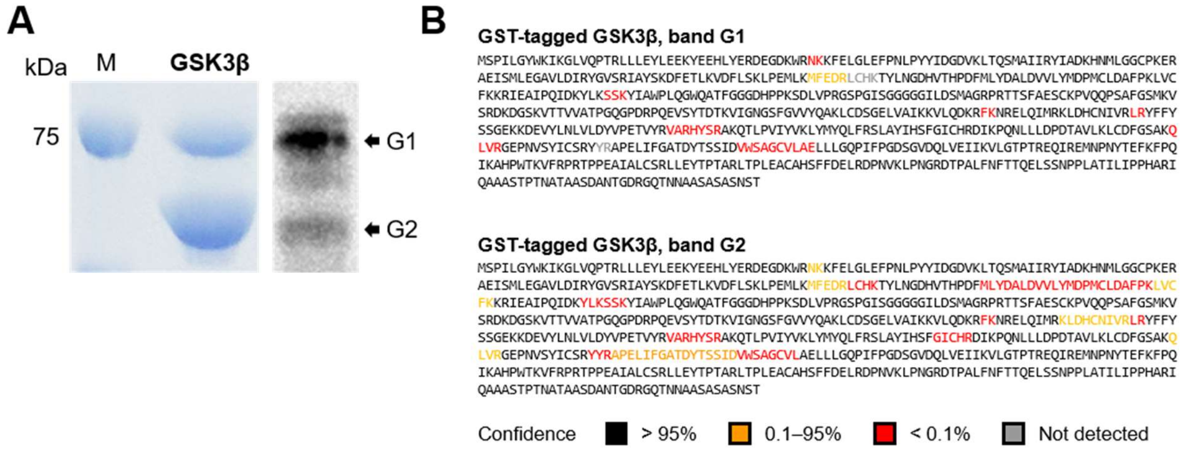


**Figure. S4. SDS-PAGE, western blot, and LC-MS analysis of purified GSK3β. (A)** SDS-PAGE (left) and Western blot (right) of purified GSK3β reveal two distinct bands (G1 and G2). **(B)** LC-MS analysis confirmed that both bands correspond to the same GSK3β sequence, verifying their common protein identity. The presence of these two bands is attributed to disulfide bonds between cysteine residues within the protein. These disulfide bonds remained incompletely reduced despite heat and reducing-agent treatment during SDS-PAGE sample preparation, leading to the observed band separation. Mass spectrometry validation confirmed the purity of GSK3β prior to use in this study.

### Cell-Based Analysis of K19 Interactions with GSK3β and 14-3-3σ

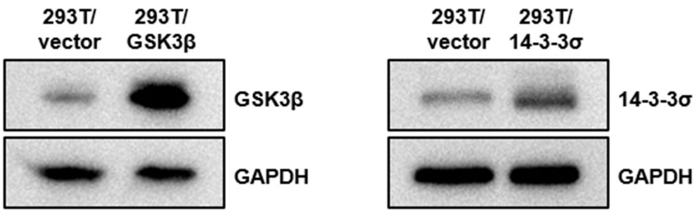


**Figure. S5. Validation of stable cell lines overexpressing GSK3β and 14-3-3σ.** Western blot analysis confirms the successful establishment of 293T stable cell lines overexpressing GSK3β (left) and 14-3-3σ (right). 293T cells transfected with the corresponding empty vector served as negative controls (293T/vector). GAPDH was used as an internal loading control.

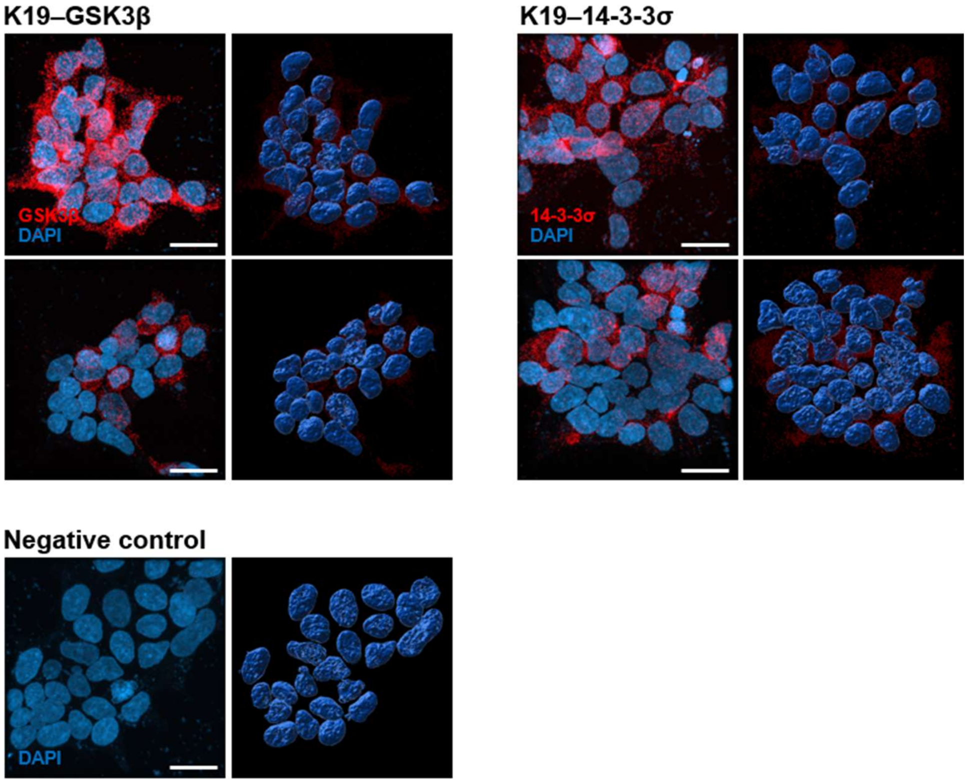


**Figure. S6.** Three-dimensional rendering of in situ protein–protein interactions detected by proximity ligation assay (PLA). Representative 3D surface-rendering images of PLA performed in the indicated 293T stable cell lines, detecting the in situ interactions of GSK3β with Keratin 19 (K19–GSK3β) and of 14-3-3σ with Keratin 19 (K19–14-3-3σ). Red signals denote positive PLA spots (individual interaction events), and blue signals denote nuclei counterstained with DAPI. The negative control (single transfection) displays only nuclear staining without red spots, confirming the specificity of the assay. Three-dimensional reconstruction was performed using IMARIS software (Bitplane, Zurich, Switzerland). Scale bars, 20 μm.
